# Processing of auditory feedback in perisylvian and insular cortex

**DOI:** 10.1101/2024.05.14.593257

**Authors:** Garret Lynn Kurteff, Alyssa M. Field, Saman Asghar, Elizabeth C. Tyler-Kabara, Dave Clarke, Howard L. Weiner, Anne E. Anderson, Andrew J. Watrous, Robert J. Buchanan, Pradeep N. Modur, Liberty S. Hamilton

## Abstract

When we speak, we not only make movements with our mouth, lips, and tongue, but we also hear the sound of our own voice. Thus, speech production in the brain involves not only controlling the movements we make, but also auditory and sensory feedback. Auditory responses are typically suppressed during speech production compared to perception, but how this manifests across space and time is unclear. Here we recorded intracranial EEG in seventeen pediatric, adolescent, and adult patients with medication-resistant epilepsy who performed a reading/listening task to investigate how other auditory responses are modulated during speech production. We identified onset and sustained responses to speech in bilateral auditory cortex, with a selective suppression of onset responses during speech production. Onset responses provide a temporal landmark during speech perception that is redundant with forward prediction during speech production. Phonological feature tuning in these “onset suppression” electrodes remained stable between perception and production. Notably, the posterior insula responded at sentence onset for both perception and production, suggesting a role in multisensory integration during feedback control.

## Introduction

A key component of speaking is the integration of ongoing sensory information from the auditory, tactile, and proprioceptive domains (Hickok, 2014; Tourville et al., 2008). When we read a sentence out loud, our brain must convert visual information into a motor program for moving our articulators (lips, jaw, tongue, larynx) to create sounds. The brain then processes these sounds as they are uttered, so the talker can hear if they sound how they expect or have made a mistake. Auditory information is processed differently during speaking compared to listening (Cogan et al., 2014; Creutzfeldt et al., 1989; Houde et al., 2002; Nourski et al., 2021; Towle et al., 2008). A prime example is speaker-induced suppression (SIS), a phenomenon in which self-generated speech generates a lower amplitude neural response than externally generated speech (Behroozmand & Larson, 2011; Flinker et al., 2010; Martikainen et al., 2005). SIS and related phenomena are components of the speech motor control system, the purpose of which is to ensure ongoing sensory feedback is in line with feedforward expectations generated prior to articulation (Guenther, 2016; Houde & Nagarajan, 2011; Tourville & Guenther, 2011). This link is established by studies that correlate the extent of cortical suppression with the accuracy of the utterance: both speech errors and subphonemic changes in utterance acoustics can result in decreased cortical suppression, indicative of a feedback control system ready to adjust the motor program in real time (Niziolek et al., 2013; Ozker et al., 2022, 2024). While feedback control has primarily been studied using noninvasive techniques with a lower signal-to-noise ratio (Chang, 2015; Houde et al., 2002; Okada et al., 2018), intracranial recordings allow for more precise investigation of this process (Chang, 2015; Hamilton, 2024; Lachaux et al., 2012; Mercier et al., 2022). This can potentially illuminate the spatiotemporal specificity of feedback suppression mechanisms like SIS. In addition, we can investigate how speech production affects other aspects of the perceptual system, such as linguistic abstraction and neural response timing.

### Organization of speech cortex during listening and speaking

Transformation of low-level acoustics into some form of intermediate linguistic representation is a necessary component of speech perception (Appelbaum, 1996). In several studies, this abstraction is organized according to place and manner of articulation, motivated by linguistic feature theory. Place of articulation describes the location of constriction in the vocal tract (e.g., a bilabial /b/ sound is produced by closing the lips). Manner of articulation, on the other hand, describes the degree of constriction and airflow through the vocal tract. Mesgarani and colleagues observed tuning of electrode populations within the superior temporal gyrus (STG) that preferentially responded to specific classes of phonological features (namely manner of speech) during passive listening (Mesgarani et al., 2014). For example, the same intracranial electrode might respond selectively to plosive phonemes such as /b/, /d/, /g/, /p/, /t/, and /k/, while not responding to fricatives such as /f/, /v/, /s/, /sh/. In more recent work, the same level of representation was observed at the single neuron level (Lakretz et al., 2021; Leonard et al., 2023). The same group later expanded on this result using a speech production task to demonstrate feature tuning changes during speech production in the motor cortex (Cheung et al., 2016). Notably, they observed that motor cortex was organized according to place of articulation during speech production, as would be expected from somatotopic representations (Bouchard et al., 2013), but organized according to manner of articulation during passive listening. However, this manuscript did not report on responses in superior temporal gyrus during speech production, nor was a direct comparison of phonological tuning made between perception and production.

A more recent insight about how the auditory system is organized comes from research on temporal response profiles in the STG (Hamilton et al., 2018). The STG contains two such profiles: first, an “onset” response region localized to posterior STG with high temporal modulation selectivity (Hullett et al., 2016) that transiently responds to the acoustic onset of a stimulus. These onset responses are useful for segmenting continuous acoustic information into discrete linguistic units, such as phrases and sentences. Second, a “sustained” response region localized to middle STG with a longer temporal integration window that does not show the same strongly adapting responses following sentence onset. Onset and sustained response profiles are a globally organizing feature of speech-responsive cortex, and responses to all phonological features are seen across both (Hamilton et al., 2018). If responses to phonological information can be modified by the acoustic context of a sound, it is possible they could also be modulated by feedback suppression during speech production. Other top-down cognitive processes can affect speech perception as well, such as expectations about upcoming stimuli evidenced in both speech production (Goregliad Fjaellingsdal et al., 2020; Lester-Smith et al., 2020; Scheerer & Jones, 2014) and speech perception (Astheimer & Sanders, 2011; Bendixen et al., 2014; Caucheteux et al., 2023). In general, auditory stimuli that are consistent with the listener’s expectations generate less of a response than inconsistent stimuli (Chao et al., 2018; Forseth et al., 2020). While consistency effects are also a component of the motor system (Gonzalez Castro et al., 2014; Shadmehr & Krakauer, 2008), the link between speaker-induced suppression and more general top-down expectation is not well established.

### Speaker induced suppression in noninvasive recordings

Recent research from our group used scalp EEG recordings to demonstrate that responses to continuous sentences are suppressed during production compared to perception of those same sentences while phonological tuning remains unchanged (Kurteff et al., 2023). However, such conclusions may be tempered by the low spatial resolution of scalp recordings, motivating the use of high-resolution intracranial stereo EEG (sEEG) recordings. When we plan to speak, the motor efference copy contains expectations about upcoming auditory feedback and may contain information about temporal/linguistic landmarks in that feedback (Levelt, 1993; Niziolek et al., 2013; Schneider et al., 2014). Onset responses, which encode the temporal landmarks of speech, may then be suppressed as a redundant processing component during speech production. This is corroborated by scalp EEG/MEG research showing that SIS occurs primarily within the N100/M100 components. That is, the N100 and M100 neural responses are suppressed during speaking as compared to playback. The N100/M100 component is an early-onset neural response that is observed at acoustic edges with high temporal modulation (Luck, 2014), making these components share characteristics with onset responses observed using invasive recordings.

### The role of the insula in speech perception and production

The use of sEEG as a recording methodology affords an additional advantage to the current study: the ability to record from deeper structures in the cortex. One such structure is the insula, a multifunctional region that is theorized to be involved in sensory, motor, and cognitive aspects of speech (Kurth et al., 2010). Recent work using sEEG reported the insula to be more active for self-generated speech when compared to externally generated speech, an opposite trend to the cortical suppression of self-generated speech observed in auditory cortex (Woolnough et al., 2019). The insula is difficult to record from using several popular neuroimaging techniques due to its placement deep in the Sylvian fissure (Chang, 2015; Remedios et al., 2009). In speech, the insula conventionally plays a role in pre-articulatory motor coordination (Dronkers, 1996). Because of the proximity of the insula to the temporal plane and hippocampus, insular coverage is rather common in sEEG epilepsy monitoring cases (Nguyen et al., 2022). We aim to expand upon the functional role of the insula in speech perception and production by directly comparing auditory feedback processing and phonological feature encoding during speaking and listening while recording from the region in high resolution.

### How do acoustic and linguistic representations change during self-produced speech?

To address how cortical suppression during speech production interacts with documented organizational phenomena during speech perception such as linguistic abstraction and onset/sustained response profiles, we used high-resolution sEEG recordings of neural activity from electrodes implanted in the cortex as part of surgical epilepsy monitoring (Guenot et al., 2001). These participants completed a dual speech production-perception task where they first read sentences aloud, then passively listened to playback of their reading to identify potential changes in local field potential recorded by the implanted electrodes. Our first goal was to identify if previously identified onset and sustained response profiles in auditory cortex (Hamilton et al., 2018) were also present during speech production. Additionally, we varied the playback condition between a consistent playback of the preceding production trial and a randomly selected playback inconsistent with the preceding trial to assess the spatial and temporal similarity of a more general perceptual expectancy effect with feedback suppression during speech production. Lastly, we investigated how linguistic feature tuning changes at individual electrodes during speech production vs. perception and how this is modulated by expectation. Our results have implications for understanding important auditory-motor interactions during natural human communication.

## Results

### Onset responses are selectively suppressed during speech production

To examine potential differences in neural processing during speech production and perception, we acquired data from 17 pediatric, adolescent, and adult participants (9F, age 16.6±6.4, range 8 to 37 years; Table S1) surgically implanted with intracranial sEEG depth electrodes and pial electrocorticography (ECoG) grids for epilepsy monitoring. These patients performed a task where they read aloud naturalistic sentence stimuli then passively listened to playback of their reading (Figure 1A). For all analyses, we extracted the high gamma analytic amplitude of the local field potentials (Lachaux et al., 2012), which has been shown to correlate with single- and multi-unit neuronal firing (Ray & Maunsell, 2011) and tracks both acoustic and phonological characteristics of speech (Mesgarani et al., 2014; Oganian et al., 2023). Based on prior work, we expected to observe strong onset and sustained responses during sentence playback (Hamilton et al., 2018, 2021), as well as sensorimotor responses during the production portions of the task that would reflect articulatory control (Bouchard & Chang, 2014; Chartier et al., 2018). Additionally, our task design allowed us to investigate the role of auditory-motor feedback during speech production by comparing neural responses to auditory feedback in real time to passive listening to an acoustically matched playback of each trial.

**Figure 1:**
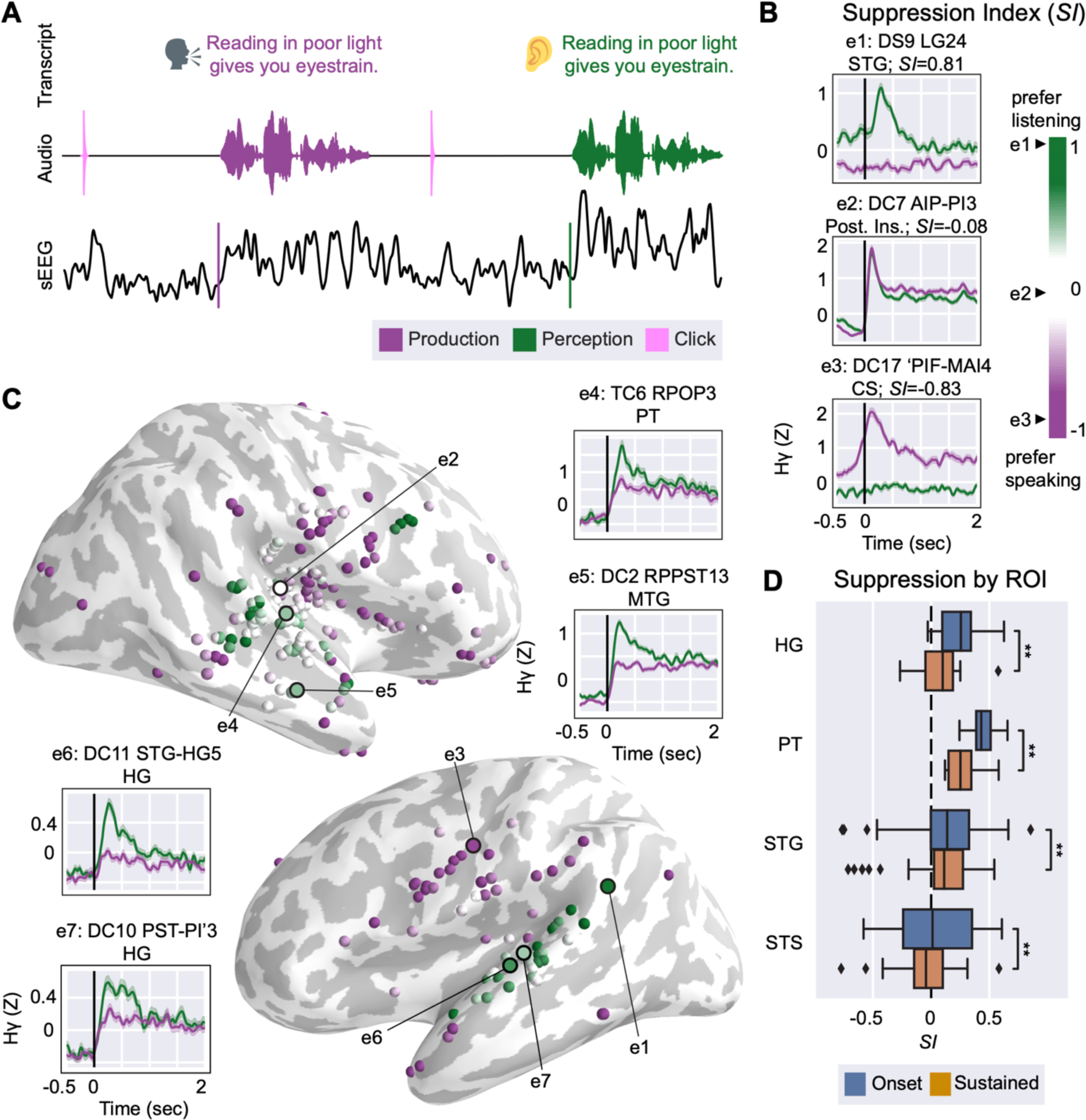
Auditory onset responses are suppressed during speech production. (A) Schematic of reading and listening task. Participants read a sentence aloud (purple) then passively listened to playback of themselves reading the sentence (green). Pink spikes in the beginning and middle of the audio waveform indicate inter-trial click tones, used as a cue and an auditory control. (B) Single-electrode plots showing different profiles of response selectivity across the cortex. Color gradient represents normalized *SI* values. A more positive *SI* indicates an electrode is more responsive to speech perception stimuli (e1) while a more negative *SI* means an electrode is more responsive to production stimuli (e3). e2 and e3 are examples of response profiles described in subsequent figures (Figures 2 and 3, respectively). Subplot titles reflect the participant ID and electrode name from the clinical montage. (C) Whole-brain and single-electrode visualizations of perception and production selectivity (*SI*). Electrodes are plotted on a template brain with an inflated cortical surface; dark gray indicates sulci while light gray indicates gyri. Single-electrode plots of high-gamma activity demonstrate suppression of onset response relative to the acoustic onset of the sentence (vertical black line). (D) Box plot of suppression index during onset (blue) and sustained (orange) time windows separated by anatomical region of interest in primary and non-primary auditory cortex. Brackets indicate significance (* = *p* < 0.05; ** = *p* < 0.01). *Abbreviations: HG: Heschl’s gyrus; PT: planum temporale; STG: superior temporal gyrus; STS: superior temporal sulcus; MTG: middle temporal gyrus; CS: central sulcus; Post. Ins.: posterior insula*.

We recorded from a total of 2044 sEEG depth electrodes implanted in perisylvian cortex and insula. This included coverage of speech responsive areas of the lateral superior temporal gyrus, but also within the depths of the superior temporal sulcus, primary auditory cortex, and surrounding regions of the temporal plane. Within- and across-subject visualizations of electrode coverage are available as supplemental figures (Figure S1, S2). To examine differences between speech perception and production on individual electrodes, we plotted event-related high gamma responses for speech perception and production trials relative to the beginning of the acoustic onset of the sentence. We identified 144 electrodes with significant responses to perceptual stimuli, 350 electrodes with significant responses to production stimuli, and 110 electrodes with significant responses to both perceptual and production stimuli (Figure 1B; bootstrap t-test, p<0.05). We quantified individual electrodes’ selectivity to speech production or perception by calculating a suppression index (*SI*, see STAR Methods).

An *SI*>0 reflects higher activity during listening compared to speaking, and *SI*<0 reflects higher activity during speaking compared to listening (Figure 1C). Single-electrode responses can be visualized on a 3D brain in an interactive webviewer at https://hamiltonlabut.github.io/kurteff2024/. We observed single electrodes with selective responses to speech perception in bilateral Heschl’s gyrus and STG (Figure 1D). 51.4% of electrodes in STG (*n* = 70) and 100% of electrodes in Heschl’s gyrus (*n* = 13) responded significantly to speech perception stimuli. Response profiles of electrodes in this region consisted of a mixture of transient onset responses and lower-amplitude sustained responses during passive listening, consistent with previous research (Hamilton et al., 2018, 2021). In primary and non-primary auditory cortex, onset responses were notably absent during speech production, while sustained responses remained relatively un-suppressed (Estimated marginal mean_onset-sustained_ *SI* = 0.153; *p* < .001). Electrodes in primary sensorimotor cortex were typically more production-selective, in line with conventional localization of sensorimotor control of speech (Bouchard et al., 2013; Guenther, 2016; Penfield & Roberts, 1959). This pattern of responses demonstrates selective suppression of onset responses during speech production in primary and secondary auditory regions of the human brain. This result supports prior research that posits onset responses play a role in temporal parcellation of speech, a process unnecessary during speech production due to the speaker’s knowledge of upcoming auditory information (Houde & Nagarajan, 2011; Tourville & Guenther, 2011).

### The posterior insula uniquely exhibits onset responses to speaking and listening

The ability of sEEG to obtain high-resolution recordings of human insula is a unique strength, as other intracranial approaches such as ECoG grids and electrocortical stimulation cannot be applied to the insula without prior dissection of the Sylvian fissure, an involved and rarely performed surgical procedure (Remedios et al., 2009; Zhang et al., 2018). Similarly, hemodynamic and lesion-based analyses may suffer from vasculature-related confounds in isolating insular responses (Hillis et al., 2004). Here we present high spatiotemporal resolution recordings from human insula and identify a functional response profile localized to this region.

While onset responses to speech perception were mostly confined to auditory cortex, a functional region of interest in the posterior insula demonstrated a different morphology of onset responses. Across participants, electrodes in the posterior insula showed robust onset responses to perceptual stimuli in similar fashion to auditory electrodes. Unlike auditory electrodes, however, posterior insular electrodes also showed robust onset responses during speech production (Figure 2D). Out of all posterior insula electrodes (*n* = 47), 23.4% responded significantly to speech perception and 31.9% responded significantly to speech production. These posterior insula onset electrodes responded similarly to stimuli regardless of whether they were spoken or heard (Figure 2). We hypothesized that such responses might reflect a relationship to articulatory motor control or somatosensory processes, which prompted us to trial a nonspeech motor control task in a subset of our participants (*n* = 6; Table S1). The purpose of this task was to determine if such “dual onset” responses were speech-specific or whether they could be elicited by simpler, speech-related movements. In this task, participants were instructed to follow instructions displayed on screen when a “go” signal was given; the instructions consisted of a variety of nonspeech oral-motor tasks taken from a typical battery used by speech-language pathologists during oral mechanism evaluations (St. Louis & Ruscello, 1981). The “go” signal contained both a visual (green circle) and an auditory cue (click), after which the participant would perform the task. Some tasks required vocalization (e.g., “say ‘aaaa’”) while others did not (e.g., “stick your tongue out”). While a few insular electrodes did exhibit responses during the speech motor control task, they were not consistently responsive to the speech motor control task except for trials that involved auditory feedback (Figure 2E). We interpret these as responses to the click sound when instructions are displayed to the participant or to the subjects’ own vocalizations rather than an index of sensorimotor activity related to the motor movements. When significance is calculated in a time window that excludes the click sound (500-100 msec post-click), only 2% of insula electrodes (*n* = 49) significantly respond to the speech motor control task. By comparison, 25.7% of sensorimotor cortex electrodes (*n* = 35) significantly responded, demonstrating that the speech motor control task was sensitive to sensorimotor activity. Additionally, posterior insular electrodes that were responsive to the speech motor control task and all dual onset insular electrodes in the main task were only active after the onset of articulation. This later response suggests that these electrodes were involved in sensory feedback processing and not direct motor control. The posterior insula region of interest was the only anatomical area in our dataset that was equally responsive to acoustic onsets during both production and perception. While electrodes with dual onset responses during speaking and listening were seen in both primary/secondary auditory areas (22.7% of dual onset electrodes) and the insula (28.8% of dual onset electrodes), electrodes with similar amplitudes for speaking and listening were most common in posterior insula (Figure 2F). In other words, while temporal electrodes did sometimes demonstrate dual onset responses, the amplitudes of these responses were larger for speech perception compared to production. We quantified this restriction of “dual onset” electrodes to posterior insula by taking the peak amplitude in the first 300 milliseconds of activity prior to sentence onset greater than 1.5 SD above the epoch mean as a measure of the onset response (Figure 2G).

**Figure 2:**
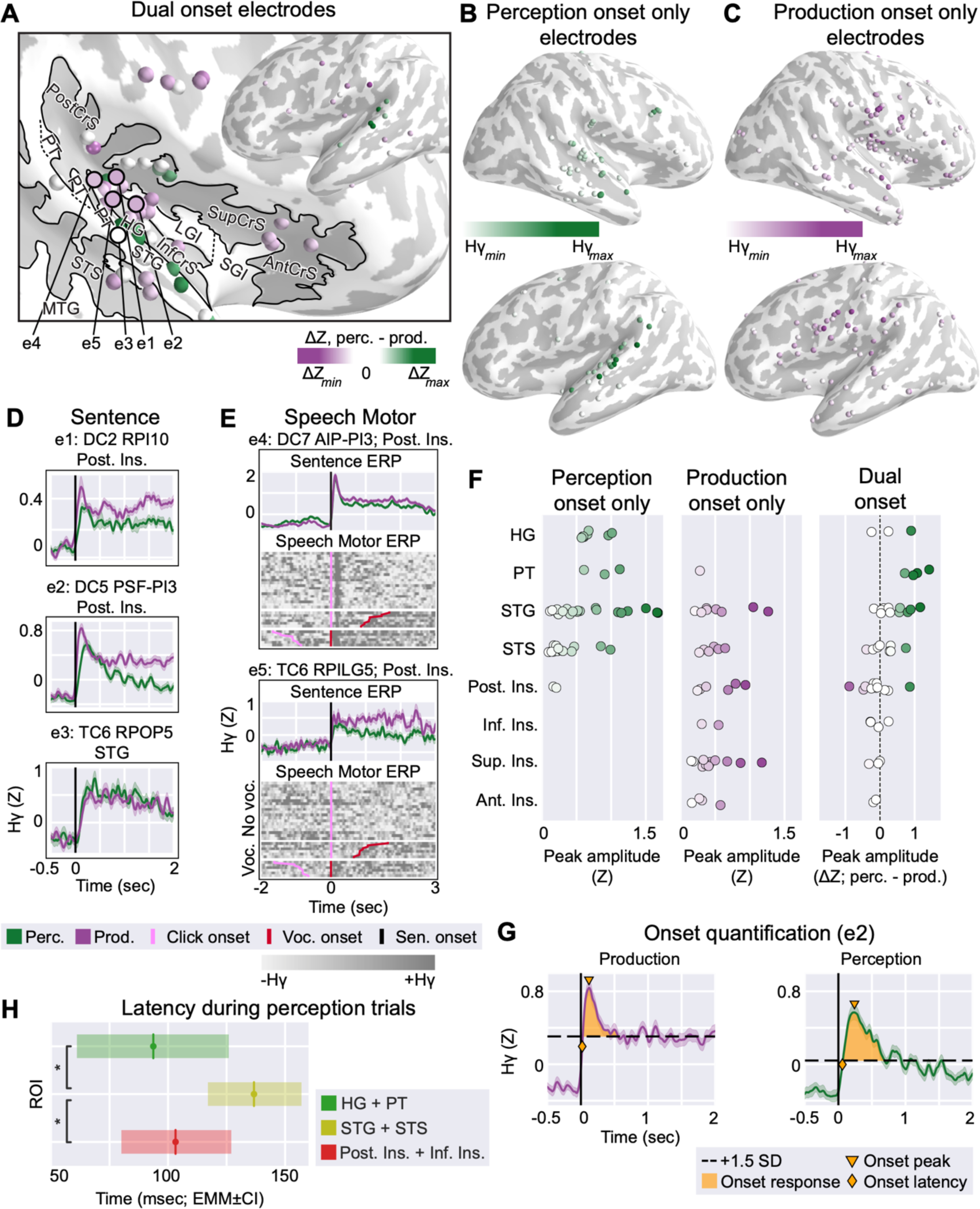
A functional region of interest in posterior insula shows onset responses to both speaking and listening. (A) Whole-brain and visualization of dual onset electrodes. Electrodes are plotted on a template brain with an inflated cortical surface; dark gray indicates sulci while light gray indicates gyri. Black outline on template brain highlights functional region of interest in posterior insula with anatomical structures labeled. Electrode color indicates the difference in Z-scored high gamma peaks during the speaking and listening conditions (ΔZ). Right hemisphere is cropped to emphasize insula ROI, while left hemisphere is shown in entirety due to lower number of electrodes. (B) Whole-brain visualization of electrodes with onset responses only during speech perception. Electrode color indicates the peak high gamma amplitude during the onset response. (C) Whole-brain visualization of electrodes with onset responses only during speech production. Electrode color indicates the peak high gamma amplitude during the onset response. (D) Single electrode activity from posterior insular electrodes highlighting dual onset responses during speech production and perception. Vertical black line indicates acoustic onset of sentence. Subplot titles reflect the participant ID, electrode name from the clinical montage, and anatomical ROI. (E) Grayscale heatmaps of single-trial electrode activity during a nonspeech motor control task, separated by no vocalization (e.g., “stick your tongue out”) and vocalization (e.g., “say ‘aaaa’”). For vocalization trials, onset of acoustic activity is visualized relative to the click accompanying the presentation of instructions (pink) and the onset of vocalization (red). (F) Strip plot showing the distribution of channel-by-channel onset response peak amplitudes separated by anatomical region of interest and whether onset responses occur only during perception (left), only during production (center), or occur during perception and production (right). Electrodes are colored according to the colormaps of (A), (B), and (C). (G) Schematic of quantification of onset response for an example electrode (e2, DC5 PSF-PI3). The first contiguous peak of activity >1.5 SD above the mean response constitutes the onset response and is shaded in orange. Peak amplitude values displayed in (B), (C) and (G) are indicated. (H) Bar plot showing the estimated marginal mean latency of the onset response in three regions of interest: auditory primary (HG + PT), auditory non-primary (STG + STS), and posterior + inferior insular. Insular onset latency is comparable to primary auditory latency. Brackets indicate significance (* = *p* < 0.05; ** = *p* < 0.01). *Abbreviations: HG: Heschl’s gyrus; STG: superior temporal gyrus; STS: superior temporal sulcus; MTG: middle temporal gyrus; Inf/Sup/Ant/Post/ CrS: inferior/superior/anterior/posterior circular sulcus of the insula; LGI: long gyrus of the insula; SGI: short gyrus of the insula; PT: planum temporale*.

The response latencies of different anatomical regions can provide a proxy for understanding how information flows from one region to another, or where in the pathway a certain response may occur. For example, our prior work showed similar latencies between the pSTG and posteromedial Heschl’s gyrus, indicating a potential parallel pathway (Hamilton et al., 2021). Here, the dual onset electrodes in posterior insula responded with comparable latency to the speech perception onset response electrodes observed in primary (HG & PT) and non-primary auditory cortex (STG & STS), in some cases responding earlier relative to sentence onset than the auditory cortex electrodes (EMM_A1_ peak latency = 93.7±16.2 msec; EMM_Aud. non-primary_ peak latency = 136.7±9.4 msec; EMM_insular_ peak latency = 103.2±11.7 msec; A1-Aud. non-primary *p* = 0.03; A1-insular *p* = 0.85; Aud. non-primary-insular *p* = 0.03; Figure 2H). This does not suggest a conventionally proposed serial cascade of information from primary auditory cortex and is instead indicative of a parallel information flow to primary auditory cortex and the posterior insula, potentially from the terminus of the ascending auditory pathway. The similar latency of posterior insular dual onset electrodes and primary auditory onset suppression electrodes alongside the tendency of posterior insular electrodes to also show low-latency onset responses during speech production leads us to speculate that the posterior insula receives a parallel thalamic input and serves as a sensory integration hub for the purposes of feedback processing during speech.

### Unsupervised identification of “onset suppression” and “dual onset” functional response profiles

Visualization of individual electrodes’ responses to the onset of perceived and produced sentences allows for manual identification of response profiles in the data but is subject to *a priori* bias by the investigators. Data driven methods such as convex non-negative matrix factorization (cNMF) allow identification of patterns in the data without access to spatial information or the acoustic content of the stimuli (Ding et al., 2010). This method was used to identify onset and sustained responses in STG (Hamilton et al., 2018). Here, we used cNMF to identify response profiles in our data in an unsupervised fashion using average evoked responses as the input to the factorization. A solution with *k* = 9 clusters explained 86% of the variance in the data (Figure 3A). We chose this threshold as increasing the number of clusters in the factorization beyond *k* = 9 resulted in redundant clusters. Similar response profiles were seen using other numbers of clusters (STAR Methods). Single-electrode responses to spoken sentences, perceived sentences, and an inter-trial click tone were used as inputs to the factorization such that responses to each of these conditions were jointly considered for defining a “cluster.” The average responses of all top-weighted electrodes within cluster for the *k* = 9 factorization is available as a supplemental figure (Figure S3). Visualization of the average response across sentences of the top-weighted electrodes within each cluster identifies two primary response profiles in correspondence with manually identified response profiles: (c1) an “onset suppression” cluster localized to bilateral STG and Heschl’s gyrus characterized by evoked responses to speech production and speech perception but an absence of onset responses during speech production; and (c2) a “dual onset” cluster localized to the posterior insula/circular sulcus characterized by evoked responses to the onset of perceived and produced sentences (Figure 3B, C). An additional cluster (c3) was localized to ventral sensorimotor cortex and showed selectivity to speech production trials, particularly prior to articulation. This cluster is located in ventral sensorimotor cortex, and likely reflects motor control of speech articulators (Bouchard et al., 2013; Breshears et al., 2015; Dichter et al., 2018).

**Figure 3.**
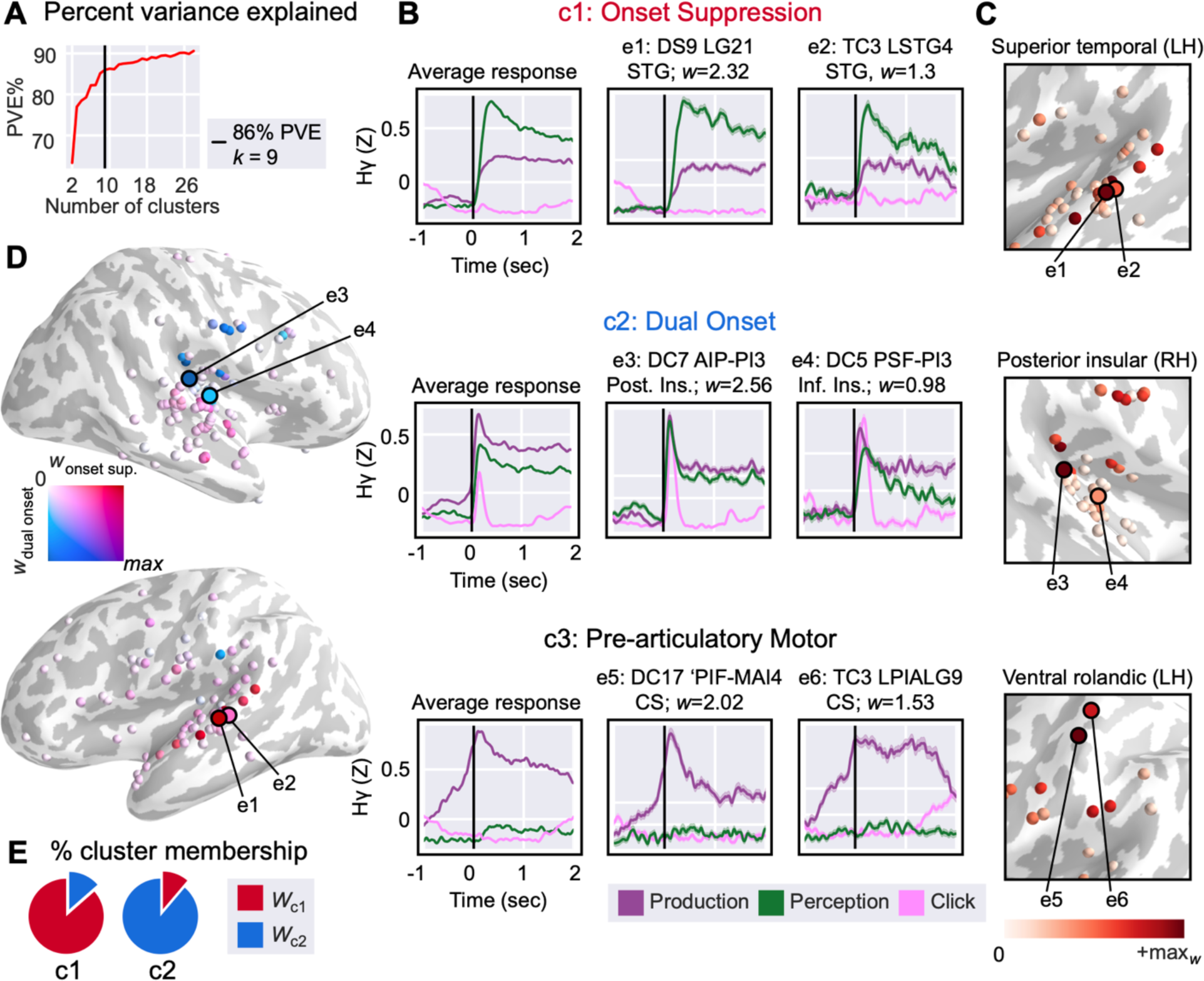
Anatomically distinct onset suppression and dual onset clusters represent a subclass of response profiles to continuous speech production and perception. (A) Percent variance explained by cNMF as a function of total number of clusters in factorization. Threshold of *k* = 9 factorization plotted as vertical black line. (B) cNMF identifies three response profiles of interest: (c1) onset suppression electrodes, characterized by a suppression of onset responses during speech production and localized to STG/HG; (c2) dual onset electrodes, characterized by the presence of onset responses during perception and production and localized to posterior insula; (c3) pre-articulatory motor electrodes, characterized by activity prior to acoustic onset of stimulus during speech production and localized to ventral sensorimotor cortex. Left: Cluster basis functions for speaking sentences (purple), listening to sentences (green), and inter-trial click (pink) for c1, c2, and c3. Center, right: Two example electrodes from the top 16 weighted electrodes. Subplot titles reflect the participant ID and electrode name from the clinical montage. (C) Cropped template brain showing top 50 weighted electrodes for individual clusters (c1, c2, c3). A darker red electrode indicates higher within-cluster weight. (D) Individual electrode contribution to dual onset and onset suppression cNMF clusters in both hemispheres. Top 50 weighted electrodes for each cluster are plotted on a template brain with an inflated cortical surface; dark gray indicates sulci while light gray indicates gyri. Red electrodes contribute more weight to the “onset suppression” cluster while blue electrodes contribute more to the “dual onset” cluster; purple electrodes contribute equally to both clusters while white electrodes contribute to neither. (E) Percent similarity of onset suppression (c1) and dual onset (c2) clusters’ top 50 electrodes. The majority of the electrode weighting across these two clusters is non-overlapping. *Abbreviations: STG: superior temporal gyrus; CS: central sulcus. Inf. Ins. = inferior insula, Post. Ins = posterior insula*.

Because the onset suppression and dual onset clusters are relatively close to each other anatomically, we quantified their functional separation by examining whether individual electrodes contributed strong weighting to both clusters. We observed that despite the spatial proximity of the clusters (which cNMF’s clustering technique would not have access to), the majority of electrodes in both onset suppression and dual onset clusters were only strongly weighted within a single cluster (Figure 3D). The top 50 electrodes of the onset suppression contributed 86.5% of their weighting to the onset suppression cluster and 13.5% to the dual onset cluster, while the top 50 electrodes of the dual onset cluster contributed 88.8% to the dual onset cluster and 11.2% to the onset suppression cluster (Figure 3E). This suggests that despite anatomical proximity, the onset responses in posterior insular electrodes are not the result of spatial spread of activity from nearby primary auditory electrodes in Heschl’s gyrus and planum temporale. Taken together, the supervised and unsupervised analyses suggest auditory feedback is processed differently by two regions in temporal and insular cortex. Auditory cortex suppresses responses to self-generated speech through attenuation of the onset response, while the posterior insula uniquely responds to onsets of auditory feedback regardless of whether the stimulus was self-generated or passively perceived.

### Response to playback consistency is a separate mechanism from suppression of onset responses

Speaker-induced suppression of self-generated auditory feedback is one example of how top-down information can influence auditory processing. In rodent studies, animals can learn to associate a particular tone frequency with self-generated movements, and motor-related auditory suppression will occur specifically for that frequency rather than unexpected frequencies that were not paired with movement (Schneider et al., 2018). Expectations about upcoming auditory feedback can also influence the outcomes of feedback perturbation tasks in humans (Lester-Smith et al., 2020; Scheerer & Jones, 2014). We were interested if other top-down expectations about the task could affect the responses of electrodes in our data and if these populations overlapped with speaker-induced suppression. To accomplish this, we separated the playback condition into blocks of consistent and inconsistent playback (Figure 4A). In the consistent playback block, participants were always played back the sentence they had just produced in the prior speaking trial. In the inconsistent playback block, participants instead were played back a randomly selected recording of a previous speaking trial. In both cases, the playback stimulus was a recording of their own voice.

**Figure 4.**
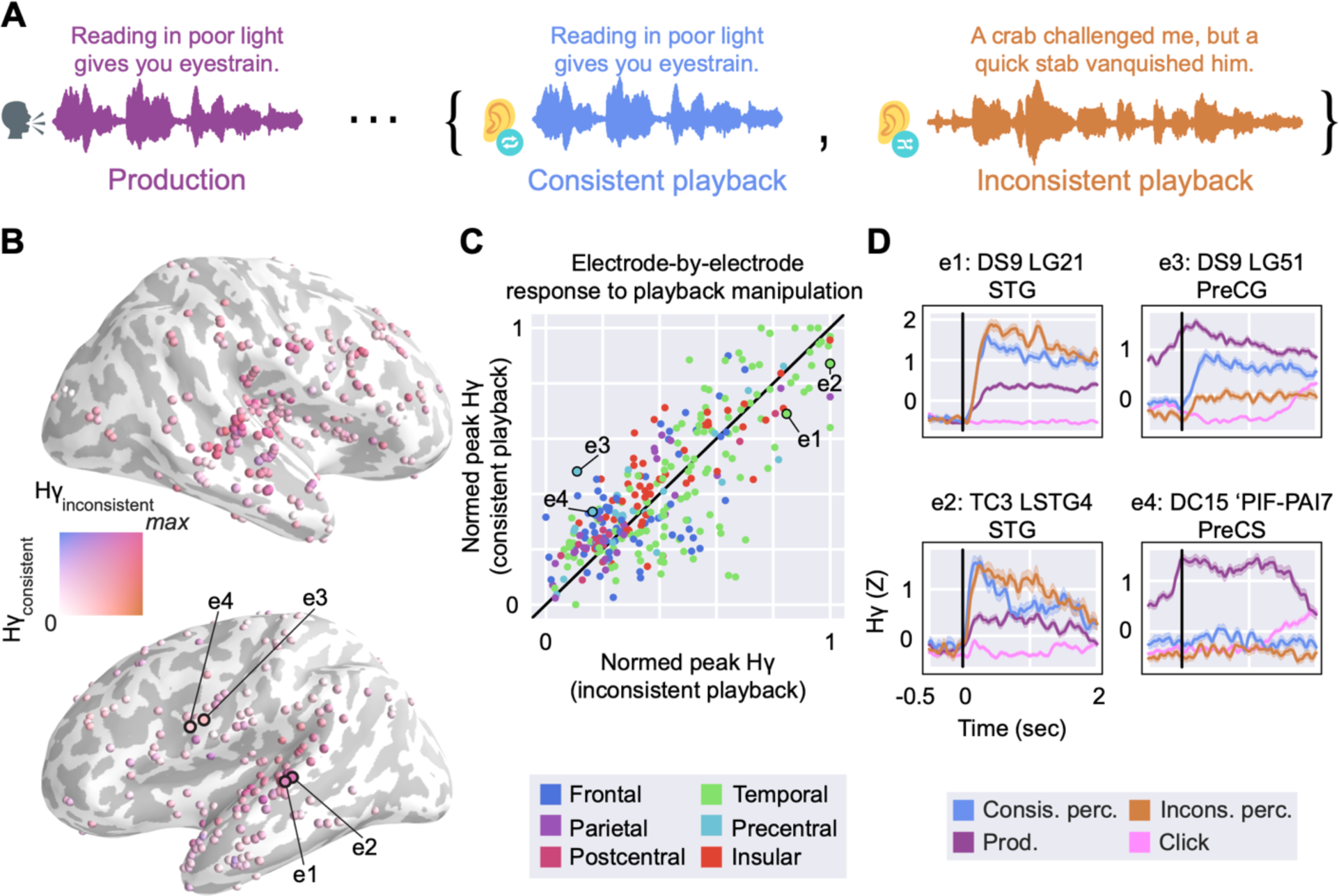
Playback consistency manipulation yields separate, weaker effects than onset suppression. (A) Task schematic showing playback consistency manipulation. Participants read a sentence aloud (purple) then passively listened to playback of that sentence (blue) or randomly selected playback of a previous trial (orange). (B) Whole-brain visualization of responsiveness to playback consistency. Electrodes are plotted on an inflated template brain; dark gray indicates sulci while light gray indicates gyri. Electrodes are colored using a 2D colormap that represents high gamma amplitude during consistent and inconsistent playback; blue indicates a response during consistent playback but not during inconsistent, orange indicates a response during inconsistent playback but not during consistent playback, pink indicates a response to both playback conditions, white indicates a response to neither. Most electrodes are pink, indicating strong responses to both conditions. Example electrodes from (D) are indicated. (C) Scatter plot of channel-by-channel peak high-gamma activity during consistent playback (Y-axis) and inconsistent playback (X-axis). Vertical black line indicates unity. Color corresponds to gross anatomical region. Example electrodes from (D) are indicated. (D) Single-electrode plots of high-gamma activity relative to sentence onset (vertical black line). Left column (e1 and e2): Electrodes in temporal cortex demonstrating a slight preference for inconsistent playback. Right column (e3 and e4): Electrodes in frontal/parietal cortex demonstrating a slight preference for consistent playback and a larger preference for speech production trials. *Abbreviations: HG: Heschl’s gyrus; STG: superior temporal gyrus; PreCS: precentral sulcus; Supramar: supramarginal gyrus*.

The majority of electrodes did not differentially respond to consistent or inconsistent playback conditions (pink-red electrodes in Figure 4B; electrodes along unity line in Figure 4C). While 45.5% of STG electrodes (*n* = 55) were significantly responsive to both consistent and inconsistent playback, only 5.5% were responsive solely during consistent playback and 0% were responsive solely during inconsistent playback. Other auditory areas showed a similar trend, including STS (both = 20.3%; consistent only = 4.3%; inconsistent only = 2.9%; *n* = 69 electrodes), posterior insula (both = 15.4%; Consistent only = 2.6%; Inconsistent only = 0%; *n* = 39 electrodes), and HG (both = 100%; Consistent only = 0%; inconsistent only = 0%; *n* = 8 electrodes). For the subset of electrodes that did differentially respond, most demonstrated a slight amplitude increase during the inconsistent playback condition that started at the time of the onset response and persisted throughout stimulus presentation (Figure 4D). Electrodes that selectively responded to inconsistent stimuli did not have an identifiable general response profile. Most electrodes that showed a preference for inconsistent playback also demonstrated onset suppression during speech production trials (e3 & e4, Figure 4D), but this suppression was far stronger than any difference between consistent and inconsistent playback. A contrast between consistent and inconsistent playback was most commonly observed in superior temporal gyrus and superior temporal sulcus. Curiously, a subset of electrodes localized to ventral sensorimotor cortex (similarly to cluster c3 presented in Figure 3B) showed an overall preference for speech production trials with pre-articulatory activity, but within the playback contrast demonstrated a preference for consistent playback (e5 & e6, Figure 4D). We interpret this finding as a speech motor region that indexes predictions of upcoming sensory content for a role in feedback control.

### Despite suppression of onset responses, phonological feature representation is suppressed but stable between perception and production

Prior work shows that circuits within the STG represent phonological feature information that is invariant to other acoustic characteristics such as pitch (Appelbaum, 1996; Mesgarani et al., 2014; Tang et al., 2017). Tuning for these phonological features is observed within both posterior onset selective areas of STG and anterior sustained regions (Hamilton et al. 2018). Here, we observed that onset responses are suppressed during speech production, which motivates investigating whether phonological feature tuning is also modulated as part of the auditory system’s differential processing of auditory information while speaking. To investigate this, we fit multivariate temporal receptive fields (mTRF) for each electrode to describe the relationship between the neural response at that electrode and selected phonological and task-level features of the stimulus (Figure 5A). We report the effectiveness of an mTRF model in predicting the neural response as the linear correlation coefficient (*r*) between a held-out validation response and the predicted response based on the model (Figure 5B, C).

**Figure 5.**
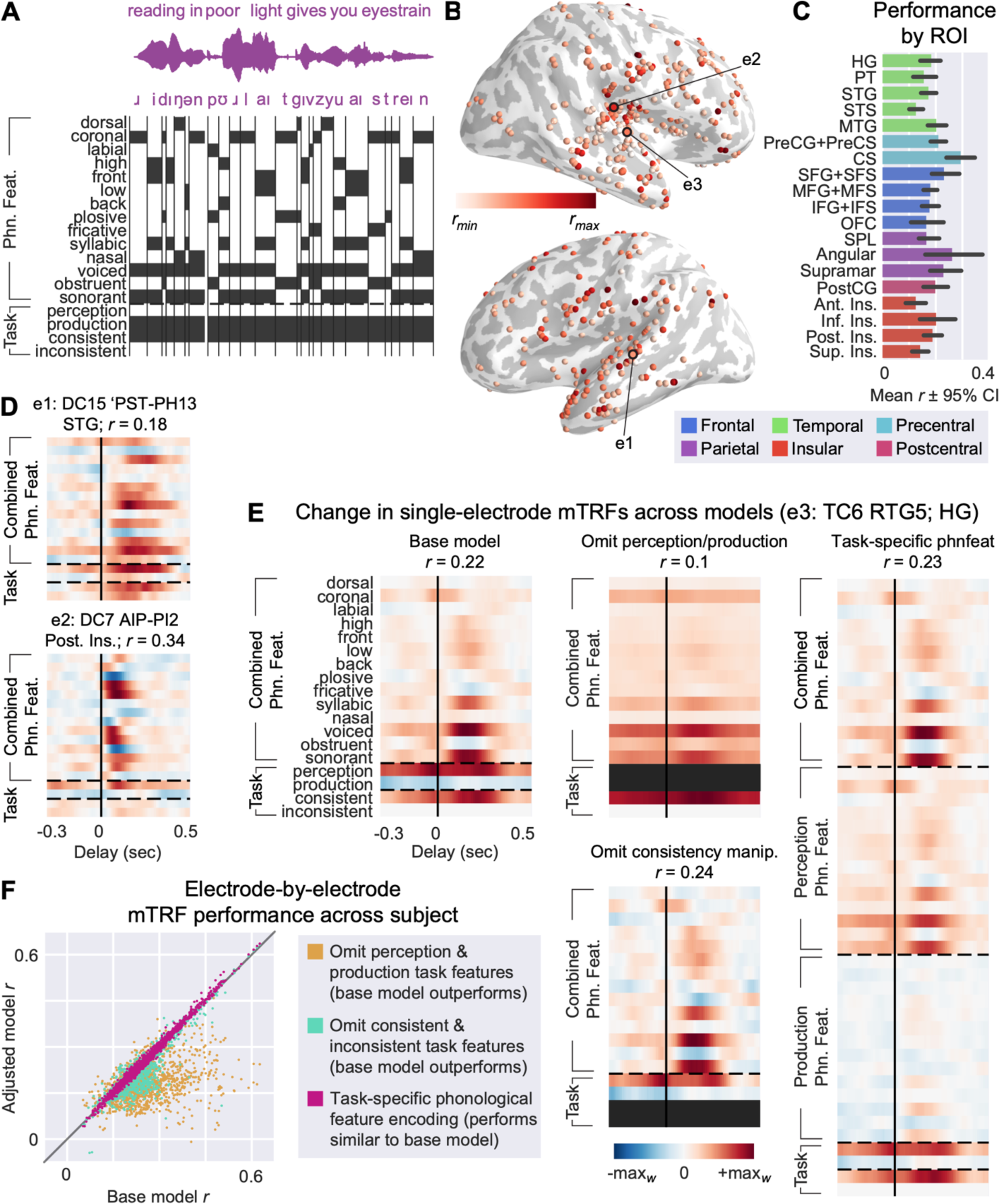
Phonological feature tuning is stable during speaking and listening across brain regions. (A) Regression schematic. Fourteen phonological features corresponding to place of articulation, manner of articulation, and presence of voicing alongside four features encoding task-specific information (i.e., whether a phoneme took place during a speaking or listening trial, the playback condition during the phoneme) were binarized sample-by-sample to form a stimulus matrix for use in temporal receptive field modeling. (B) Model performance as measured by the linear correlation coefficient (*r*) between the model’s prediction of the held-out sEEG and the actual response plotted at an individual electrode level on an inflated template brain; dark gray indicates sulci while light gray indicates gyri. Example electrodes from (D) and (E) are indicated. (C) Model performance by region of interest. Color corresponds to gross anatomical region. (D) Temporal receptive fields of two example electrodes in temporal and insular cortex. (E) Temporal receptive fields of an example electrode for the four models presented in (F). (F) Scatter plot of channel-by-channel linear correlation coefficients (*r*) colored by model comparison. The X-axis shows performance for the “base” model whose schematic is presented in (A). The Y-axis for each scatterplot shows performance for a modified version of the base model: task features encoding production and perception were removed from the model (yellow); task features encoding consistent and inconsistent playback conditions were removed from the model (cyan); phonological features were separated into production-specific, perception-specific, and combined spaces (magenta). *Abbreviations: HG: Heschl’s gyrus; PT: planum temporale; STG/S: superior temporal gyrus/sulcus; MTG/S: middle temporal gyrus/sulcus; PreCG/S: precentral gyrus/sulcus; CS: central sulcus; SFG/S: superior frontal gyrus/sulcus; MFG/S: middle frontal gyrus/sulcus; IFG/S: inferior frontal gyrus/sulcus; OFC: orbitofrontal cortex; SPL: superior parietal lobule; PostCG: postcentral gyrus; Ant./Post./Sup./Inf. Ins.: anterior/posterior/superior/inferior insula*.

Onset suppression electrodes in auditory cortex and dual onset electrodes in the posterior insula were both well modeled using this approach (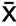*r*_onset suppression electrodes_ = 0.17±0.08; 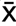*r*_dual onset electrodes_ = 0.16±0.11, range -0.25 to 0.64; Figure 5D). Within both response profiles, single electrodes exhibited a diversity of preferences to various combinations of phonological features, mirroring previous results showing distributed phonological feature tuning in auditory cortex (Berezutskaya et al., 2017; Hamilton et al., 2018, 2021; Mesgarani et al., 2014; Oganian & Chang, 2019). Of note, posterior and inferior insula electrodes were strongly phonologically tuned, with a short temporal response profile as was seen in our prior latency analysis. Dual onset and onset suppression electrodes differed from purely production-selective electrodes in this way, as most production-selective electrodes qualitatively did not demonstrate robust phonological feature tuning. Instead, most of the variance in the mTRF instead was explainable by global task-related stimulus features (i.e., whether a sound occurred during a production or a perception trial).

To directly compare phonological feature representations during perception and production, we used variance partitioning techniques to omit or include specific stimulus features in our model. In this way, the stimulus matrix serves as a hypothesis about what stimulus characteristics will be important in modeling the neural response. Adding or removing individual stimulus characteristics and observing differences (or lack thereof) in model performance serves as a causal technique for assessing the importance of a stimulus characteristic to the variance of an electrode’s response (Ivanova et al., 2021). In the base model, we included 14 phonological features and 4 task-related features. We first expanded the specificity of phonological feature tuning in our stimulus matrix by separating the phonological feature space into whether the phonemes in question occurred during perception or production (called the “task-specific” model). If phonological feature tuning differed during speech production, model performance should increase when modeling perceived vs. produced phonological features separately. However, we saw no significant increase in model performance when expanding the model in this way (Figure 5F, pink points). Despite no gross difference in model performance, inspection of individual electrodes’ receptive fields shows a suppression in the weights for production-specific phonological feature tuning (Figure 5E, far right). Still, this difference was not statistically significant, thus favoring the simpler “base” model (EMM_base - task-specific phnfeat_ Δ*r* = -0.002, *p* = 0.12, *d* = -0.05).

Removal of the playback consistency information from the task-specific portion of the stimulus matrix similarly does affect model performance; however, the effect is quantitatively weak (EMM_base - omit consistent/inconsistent_ Δ*r* = 0.01, *p* < .001, *d* = 0.02). On the other hand, removing information about the contrast of perception and production trials entirely from the model more drastically impairs model performance (EMM_base - omit perception/production_ Δ*r* = 0.07, *p* < .001, *d* = .93). Upon inspection, the regions exhibiting the largest decline in encoding performance with the omission of the perception-production contrast are frontal production-responsive regions and temporal onset suppression regions, whereas insular electrodes did not see as steep a decline in performance. This suggests that differences in encoding during speech production and perception are the primary explanation of variance in our models. Ultimately, despite onset suppression seen during speech production, higher-order linguistic representations such as phonological features appear to be stable during speech perception and production.

Taken together, these results provide an expanded perspective on how auditory areas of the brain differentially process sensory information during speech production and perception. Transient responses to acoustic onsets in primary and higher order auditory areas are suppressed during speech production, whereas responses of these regions not at acoustic onset remain relatively stable between perception and production. This onset suppression can be seen in the neural time series and is also reflected in the encoding of linguistic information in temporal receptive field models. It is thus possible that the onset response functions as a stimulus orientation mechanism rather than a higher-order aspect of the perceptual system such as phonological encoding. While expectations about the linguistic content of upcoming auditory playback can influence response profiles, the mechanism appears separate from the suppression of onset responses and is a relatively weak effect by comparison. Lastly, these results provide a unique perspective on the role of the posterior insula during speaking and listening, characterized by its rapid responses to speech production and perception stimuli and phonological tuning without the suppression observed during speech production in nearby temporal areas.

## Discussion

We used a sentence reading and playback task that allowed us to compare mechanisms of auditory perception and production while controlling for stimulus acoustics. The primary objective was to assess spatiotemporal differences in previously identified onset and sustained response profiles in the auditory cortex (Hamilton et al., 2018) and phonological feature encoding (Mesgarani et al., 2014) during speech production. Using sEEG has the distinct advantage of penetrating into deeper structures inside the Sylvian fissure, such as the insula and Heschl’s gyrus (Chang, 2015). In temporal cortex, proximal to where onset responses have been previously identified using surface electrocorticography (Hamilton et al., 2018), we observed a selective suppression of transient responses to sentence onset during speech production, whereas sustained responses remained relatively unchanged between speech perception and production. The timing of the suppressed onset responses is roughly aligned with scalp-based studies of speaker-induced suppression that posit early components (N1 for EEG, M1 for MEG) as biomarkers of speaker-induced suppression (Hawco et al., 2009; Heinks-Maldonado et al., 2006; Kurteff et al., 2023; Martikainen et al., 2005). While we do not claim the onset responses observed in our study and others to be equivalent to N/M100, there is a parallel to be drawn between the temporal characteristics of our suppressed cortical activity and the deep literature on suppression of these components during speech production in noninvasive studies. In the original onset and sustained response profile paper (Hamilton et al., 2018), the authors theorized that onset responses may serve a role as an auditory cue detection mechanism based on their utility to detect phrase and sentence boundaries in a decoder framework. Novel stimulus orienting responses have been localized to middle and superior temporal gyrus, which overlaps with the functional region of interest for onset responses (Friedman et al., 2009). These findings are in line with the absence of onset responses during speech production, as auditory orientation mechanisms during speech perception are not necessary to the same extent during speech production due to the presence of a robust forward model of upcoming sensory information (i.e., efference copy) generated as part of the speech planning process (Houde & Chang, 2015; Tourville & Guenther, 2011). A notable difference between the original reporting of onset and sustained response profiles in Hamilton et al., 2018 and the current study is that many of the electrodes reported in our analysis showed a mixture of onset and sustained response profiles, whereas the original paper posits a more stark contrast in the response profiles. This could be due to differences in coverage between the sEEG depth electrodes used here and the pial ECoG grids used in the original study, as the onset response profile was reported to be localized to a relatively small portion of dorsal-posterior STG. Many of onset electrodes were recorded from within STS or other parts of STG; therefore, the activity recorded at those electrodes may represent a mixture of onset and sustained response, which explains why both would show up in the averaged waveform. Mixed onset-and-sustained responses have been previously reported primarily in HG/PT in a study using ECoG grids covering the temporal plane (Hamilton et al., 2021); our use of sEEG depths may be providing greater coverage of these intra-Sylvian structures. Alternatively, the mixed onset-sustained responses we see in our data may be a mixture of the onset region with the posterior subset of sustained electrodes reported in the original paper. We did observe solely onset-responsive and solely sustained-responsive electrodes (in line with the original paper), but a majority of the onset suppression response profile described in this study consisted of a mixture of onset and sustained responses at the single electrode level. Responses to the inter-trial click tone observed at some electrodes are another example of pure onset response electrodes in these data.

The suppression of onset responses in temporal cortex did not impact the structure of phonological feature representations for these electrodes. Phonological feature tuning has been demonstrated previously during speech production, but the analysis focused primarily on motor cortex and not a direct comparison to the representations present in temporal cortex during speech perception (Cheung et al., 2016). In the present study, an encoding model capable of differentially encoding phonological features during speech perception and production did not outperform a model only capable of encoding phonological features identically during perception and production, demonstrating that differences in encoding performance during speech production are not due to changes in the phonological feature tuning of individual electrodes. In other words, an electrode that encodes plosive voiced obstruents (like /b/, /g/, /d/) during speech perception will still encode plosive voiced obstruents during speaker-induced suppression, but the amplitude of the response is reduced during speaking. This is consistent with similar research in scalp EEG conducted by our group (Kurteff et al., 2023) and supports the confinement of cortical suppression during speech production strictly to lower-level sensory components of the auditory system. This is also in line with previous literature showing the degree of suppression observed at an individual utterance is dependent on that utterance’s adherence to a sensory goal (Niziolek et al., 2013).

In our analysis, the posterior insula served as a unique functional region in processing auditory feedback during speech production and perception. Unlike temporal cortex, onset responses were not suppressed during speech production in posterior insula; the region instead exhibited “dual onset” responses during speech production and perception. A large portion of the research on the human insula’s involvement in speech and language comes from lesion and functional imaging studies that posit a preparatory motor role for the insula in speech (Ackermann & Riecker, 2004; Dronkers, 1996; Mandelli et al., 2014). However, these studies prescribe this role to the anterior insula, whereas our findings are constrained to posterior insula, and the insula is far from anatomically or functionally homogenous (Kurth et al., 2010; Quabs et al., 2022; Zhang et al., 2018). A meta-analysis of the functional role of human insula parcellated the lobe into four primary zones: social-emotional, cognitive, sensorimotor, and olfactory-gustatory (Kurth et al., 2010). As speech production involves sensorimotor and cognitive processes, even speech cannot be constrained to one functional region of the insula. Cytoarchitectonically, the human insula consists of eleven distinct regions which can be grossly clustered into three zones: a dorsal-posterior granular-dysgranular zone, a ventral-middle-posterior agranular-dysgranular zone, and a dorsal-anterior granular zone (Quabs et al., 2022). Based on the general organizational principles of these articles, the dual onset responses we observed in the posterior insula overlap with functional regions of interest for somatosensory, motor, speech, and interoceptive function, and with the dorsal-posterior and ventral-middle-posterior cytoarchitectonic zones. The posterior insula responses we report in this study are purely post-articulatory, indicating a role in auditory feedback monitoring rather than a preparatory motor role. This is corroborated by a recent study that identified an auditory region in dorsal-posterior insula through intraoperative electrocortical stimulation (Zhang et al., 2018), whereby stimulation to posterior insula resulted in auditory hallucinations. Several studies using animal models, including nonhuman primates, have also identified an auditory field in the posterior insula (Linke & Schwegler, 2000; Remedios et al., 2009; Rodgers et al., 2008). While this insular auditory field does receive input from primary and secondary auditory areas, it also receives direct parallel input from the auditory thalamus, evidenced in part by pure-tone responses in the insular auditory field sometimes having a lower response latency than the primary auditory cortex (Jankowski et al., 2023; Sawatari et al., 2011; Takemoto et al., 2014). Our own results parallel animal models, as we observed faster (or equivalently fast) responses to auditory playback stimuli in the posterior insula compared to primary (HG, PT) and higher order (STG, STS) auditory areas. Thus, this study corroborates parallel auditory pathways between auditory cortex and posterior insula but in the human brain and with more complex auditory stimuli than pure tones. We also expand upon animal models by showing responses to auditory feedback in insula are also present during speech production.

While posterior insula and HG are neighboring anatomical structures, we do not believe our posterior insula responses to be simply miscategorized HG activity due to the distinction between how HG and posterior insula respectively suppress or do not suppress auditory feedback during speech production. This is corroborated by the functional separation of cluster weights in our cNMF analysis between “onset suppression” and “dual onset” electrodes, alongside the fact that the high gamma LFP we report on has lower spatial spread than other frequency bands (Muller et al., 2016). Our data are by no means the first to report *in vivo* recordings of the human insula’s responses to speech perception and production: Woolnough et al., 2019 also reported post-articulatory activity in the human insula during speech production and perception. Our insular results are distinct from this study in several ways. First, the authors dichotomize the posterior insula with STG, reporting that posterior insula is more active for self-generated speech “opposite of STG.” However, our dual onset response electrodes in the posterior insula are equivalently responsive to speech perception and production stimuli, with only a small non-significant preference for speech production. Second, the responses reported in this paper differ in magnitude between STG and the posterior insula, with task-evoked activity in STG increasing ∼200% in broadband gamma activity from baseline, while posterior insula showed only ∼50% increase in activity from baseline. In our results, temporal and insular evoked activity are similar in magnitude. Third, the authors used separate tasks with distinct stimuli to compare perception and production, while we generated perceptual stimuli from individual participants’ own utterances, allowing us to control for temporal and spectral characteristics of the stimuli and more directly compare speech perception with production within the posterior insula for the same stimulus. We interpret the posterior insula’s role in speech production as a hub for integrating the multiple modalities of sensory feedback (e.g., auditory, tactile, proprioceptive) available during speech production for the purposes of speech monitoring, based in part on previous work establishing the insula’s role in multisensory integration (Kurth et al., 2010). Diffusion tensor imaging reveals that the posterior insula in particular is characterized by strong connectivity to auditory, sensorimotor, and visual cortices, supporting such a role (Zhang et al., 2018). Our research motivates further investigation of the role of the posterior insula in feedback control of speech production.

While the primary focus of this study was to describe differences in auditory feedback processing during perception and production, we were motivated to include a consistency manipulation within our speech perception condition by several findings. Behaviorally, participants’ habituation to the task can affect results: inconsistent perturbations of feedback during a feedback perturbation task elicit larger corrective responses than consistent, expected perturbations (Lester-Smith et al., 2020). The importance of predicting upcoming sensory consequences is visible in the neural data as well: unpredicted auditory stimuli result in suppression of scalp EEG components for self-generated speech in pitch perturbation studies (Scheerer & Jones, 2014) as well as the speech of others in a turn-taking sentence production task (Goregliad Fjaellingsdal et al., 2020). We sought to delineate whether onset responses were an important component of specifically speech perception or involved in a more general predictive processing system. While we did observe that presenting auditory playback in a randomized, inconsistent fashion resulted in a greater response amplitude for some onset suppression electrodes in auditory cortex, this finding did not hold true for most onset suppression electrodes in our data. This leads us to believe that the suppression of onset responses is not a byproduct of general expectancy mechanisms modulating the speech perception system, but rather a dedicated component of auditory processing for orienting to novel stimuli. Cortical suppression of self-generated sounds is likely a fundamental component of the sensorimotor system, as neural responses to tones paired with non-speech movements are attenuated relative to unpaired tones in mice and in humans (Martikainen et al., 2005; Schneider et al., 2018). With cNMF, we identified a cluster in ventral sensorimotor cortex that was more active for speech production, but within the consistent/inconsistent playback split, preferred consistent playback. We interpret this response profile as indicative of feedback enhancement for the purposes of speech motor control during speech production. This playback consistency manipulation was also included in a recently published EEG version of this task (Kurteff et al., 2023), but the results of the manipulation were inconclusive. In that EEG study, however, we did see cortical suppression at sentence onset, so perhaps the lack of a result for the consistency manipulation is a mixture of the relatively smaller effect size of the consistency manipulation and the lower signal-to-noise ratio of scalp EEG recordings in comparison to intracranial EEG.

Because our dataset uses sEEG depth electrodes, we were able to record from a wide array of cortical and subcortical areas impractical to cover with ECoG grids. As a result, there were several interesting trends observed within single subjects that were not robust enough to report upon earlier but do warrant a more speculative discussion. Occipital coverage was generally limited for this study, but one subject (DC7) had three electrodes in the right lateral occipital cortex that strongly preferentially responded to speech production trials and to click responses (DC7 PT-MT15 *p*_production_ = 0.01; *p*_perception_ = 0.9). We identified this area using our unsupervised clustering analysis: cNMF identified a cluster selective to clicks and speech production localized to the occipital lobe (Figure S3, cluster 6). We interpret this as a byproduct of our task design, as text was displayed during speech production trials (the sentence to be read aloud) but not during perception trials. The between-peak duration of the bimodal click response observed in the cNMF cluster is ∼1000 msec, which corresponds with the amount of time a fixation cross was displayed at the beginning of each trial (see STAR Methods). Based on this information, we conclude these occipital electrodes for DC7 are encoding visual scene changes between fixation cross and text display, but we advise caution in generalizing this to a functional localization as we only observed this trend in a single subject. In a separate single subject (DC5), we observed electrodes in the right inferior frontal sulcus (just dorsal of pars triangularis of the inferior frontal gyrus) that responded selectively to speech perception and inter-trial click tones (DC5 AMF-AI4 *p*_production_ = 0.31; *p*_perception_ < .001). Unlike onset suppression electrodes in auditory cortex, these electrodes were silent during speech production for onset and sustained responses. The amplitude of production responses increased as the depth progressed laterally towards pars triangularis, but the final electrode of the depth still had a (barely) non-significant response to speech production trials (DC5 AMF-AI8 *p*_production_ = 0.06; *p*_perception_ = 0.45). Unlike the occipital electrodes described above, the inferior frontal perception-selective electrodes of DC5 did not emerge as a functional region in our unsupervised clustering analysis and were interspersed with other perception-selective electrodes from other subjects localized to PT and HG (Figure S3, cluster 7). While the convention of inferior frontal cortex being monolithically a speech production region is increasingly being challenged in contemporary research (Fedorenko & Blank, 2020; Flinker et al., 2015; Hickok et al., 2023; Tremblay & Dick, 2016), the confinement of our perception-selective electrodes in this region to a single subject gives us hesitation to weigh in on this topic.

Overall, this project gives clarity to both the differential processing of the auditory system during speech production and the functional role of onset responses as a temporal landmark detection mechanism through high-resolution intracranial recordings of a naturalistic speech production and perception task. To be specific, the suppression of onset responses during speech production lends to the hypothesis that onset responses are an orientational mechanism. Feedforward expectations about upcoming sensory feedback during speech production would nullify the need for temporal landmark detection to the same extent necessary during speech perception, where expectations about incoming sensory content are much less precise. This raises questions about the function of onset responses in populations with disordered feedforward/feedback control systems, such as apraxia of speech (Jacks & Haley, 2015), schizophrenia (Heinks-Maldonado et al., 2007), and stuttering (Max & Daliri, 2019; Toyomura et al., 2020). The presence or absence of onset responses having no effect on the structure of phonological feature representations also supports this hypothesis, as linguistic abstraction is a higher-level perceptual mechanism that need not be implicated in lower-level processing of the auditory system. In future studies, we would like to further investigate the role of onset responses in less typical speech production. Just as self-generated speech is less suppressed during errors (Ozker et al., 2022, 2024) and less canonical utterances (Niziolek et al., 2013), the landmark detection services of the onset response may be more necessary in these contexts, leading to a reduced suppression of the onset response. Future research should also aim to better dissociate onset responses from expectancy effects observed in feedback perturbation tasks, which are similar in terms of spatial and temporal profile to onset responses in our data due to the limitations of naturalistic study design, yet we speculate mechanistically different than onset responses. Our findings support a functional network between the lateral temporal lobe, insula, and motor cortex to support natural communication. The differential responses of the speech-regions of STG and insula support the role of the posterior insula in auditory feedback control during speaking.

## Acknowledgements

The authors would like to thank the patients at Dell Children’s Medical Center, Texas Children’s Medical Center, and Dell Seton Medical Center for volunteering time during their hospital stay to participate in this research. We thank the members of the clinical team at Dell Children’s that assisted in data collection and/or patient referral: Timothy George, MD; Winson Ho, MD; Nancy Nussbaum, PhD; Rosario DeLeon, PhD; William Andy Schraegle, PhD; Fred Perkins, MD; Karen Keough, MD; Aaron Cardon, MD; Karen Skjei, MD; Teresa Ontiveros, RN, MSN; Cassidy Wink, RN; and Bethany Hepokoski, RN. We thank Maansi Desai, PhD for her assistance with data collection and manuscript edits. Lastly, we thank Stephanie Ries, PhD; Maya Henry, PhD, CCC-SLP; Rosemary A. Lester-Smith PhD, CCC-SLP; and Jun Wang, PhD for their feedback on early manuscript drafts. Funding was provided by the National Institutes of Health National Institute on Deafness and Other Communication Disorders (1R01-DC018579, to L.S.H.) and by a William Orr Dingwall Foundation Dissertation Fellowship (to G.L.K.).

## Author contributions

Conceptualization: G.L.K., L.S.H.; methodology: G.L.K., L.S.H.; software: G.L.K., L.S.H.; formal analysis: G.L.K., L.S.H.; investigation: all authors; data curation: G.L.K., S.A., A.F., and L.S.H.; writing – original draft: G.L.K., L.S.H.; writing – review and editing: all authors; visualization: G.L.K., L.S.H.; supervision: G.L.K., L.S.H.; project administration: G.L.K., L.S.H.; funding acquisition: L.S.H.

## STAR Methods

### Key resources table

| REAGENT or RESOURCE | SOURCE | IDENTIFIER |
| --- | --- | --- |
| Software and algorithms |  |  |
| Python 3.9.7 | python.org | N/A |
| MNE 1.1.1 | Gramfort et al. (Gramfort et al., 2013) | https://doi.org/10.1016/j.neuroimage.2013.10.027 |
| MATLAB r2021b | mathworks.com | N/A |
| R 4.2.1 | r-project.org | N/A |
| Custom code and data | This paper | GitHub for code:<br>https://github.com/HamiltonLabUT/kurteff2024_code<br>Data will be made available through contact to the lead author |
| Imaging pipeline for stereotactic localization of electrodes | Hamilton et al. (Hamilton et al., 2017) | https://doi.org/10.3389/fninf.2017.00062 |
| Browser-based electrode viewer | This paper | https://hamiltonlabut.github.io/kurteff2024/ |
| Other |  |  |
| Human patient participants recruited from Dell Children's Medical Center, Dell Seton Medical Center, and Texas Children's Hospital (see Table S1) | This paper | N/A |

## Resource availability

### Lead contact

Further information and requests for resources and reagents should be directed to and will be fulfilled by the lead contact, Liberty S. Hamilton.

### Materials availability

This study did not generate new unique reagents.

### Data and code availability

- The neural data reported in this study cannot be deposited in a public repository because they could compromise research participant privacy and consent. To request access, contact the lead contact.
- All original code has been deposited at GitHub and is publicly available as of the date of publication. URLs are listed in the key resources table.
- Any additional information required to reanalyze the data reported in this paper is available from the lead contact upon request.

### Experimental model and subject details

17 individuals (sex: 9F; age: 16.6±6.4, range 8-37; race/ethnicity: 8 Hispanic/Latino, 6 White, 1 Asian, 2 multi-racial) undergoing intracranial monitoring of seizure activity via stereoelectroencephalography (sEEG) for medically intractable epilepsy were recruited from three hospitals: Dell Children’s Medical Center in Austin, Texas (*n* = 13); Texas Children’s Hospital in Houston (*n* = 3), Texas; and Dell Seton Medical Center in Austin, Texas (*n* = 1). Demographic and relevant clinical information is provided in Table S1. Participants (and for minors, their guardians) received informed consent and provided written consent for participation in the study. All experimental procedures were approved by the Institutional Review Board at the University of Texas at Austin.

## Method details

### Neural data acquisition

Intracranial sEEG and ECoG data from a total of 2044 electrodes across subjects were recorded continuously via the epilepsy monitoring teams using a Natus Quantum headbox (Natus Medical Incorporated, San Carlos, CA, USA). At Texas Children’s Hospital, sEEG depths (AdTech Spencer Probe Depth electrodes, 5mm spacing, 0.86mm diameter, 4-16 contacts per device), strip electrodes (AdTech) and grids (AdTech custom order, 5mm spacing, 8×8 contacts per device) were implanted in the brain by the neurosurgeon in brain areas that are determined via clinical need. At Dell Children’s Medical Center and Dell Seton Medical Center, sEEG depths (PMT Depthalon, 0.8mm diameter, 3.5mm spacing, 4-16 contacts per device) were used. A TDT S-Box splitter was used at Dell Children’s Medical Center to connect the data stream to a TDT PZ5 amplifier, which then recorded the local field potential from the sEEG electrodes onto a research computer running TDT Synapse via a TDT RZ2 digital signal processor (Tucker Davis Technologies, Alachua, FL, USA). Speaker (perceived) and microphone (produced) audio were also recorded via RZ2 at 22 kHZ to circumvent downsampling of audio by the clinical recording system. At the other two recording locations, use of a dedicated research recording system was not possible due to clinical constraints; instead, the auditory stimuli from the iPad were recorded directly on the clinical system using an audio splitter cable. Simultaneous high-resolution audio was recorded for both speaking and playback using an external microphone and a second splitter cable from the iPad both plugged into a MOTU M4 USB audio interface (MOTU, Cambridge, MA, USA) plugged into the research computer running Audacity recording software. After the recording session, a match filter was used to synchronize high-resolution audio from the external recording system to the neural data recorded on the clinical system (Turin, 1960). Intracranial data were recorded at 3 kHz and downsampled to 512 Hz before analysis for all sites.

### Data preprocessing

Data were preprocessed offline using a combination of custom MATLAB scripts and custom Python scripts built off the MNE-python software package (Gramfort et al., 2013). First, data were notch filtered at 60/120/180 Hz to remove line noise, then bad channels were manually inspected and rejected. Next, a common average reference was applied across all non-bad channels. The high gamma analytic amplitude response (Lachaux et al., 2012), which has been shown to strongly correlate with speech (Kunii et al., 2013) and serves as a proxy for multi-unit neuronal firing (Ray & Maunsell, 2011), was extracted via Hilbert transform (8 bands, log spaced, Gaussian kernel, 70-150 Hz). Lastly, the 8-band Hilbert transform response was Z-scored relative to the mean activity of the individual recording block. All preprocessing and subsequent analyses were performed on a research computer with the following specifications: Ubuntu 20.04, AMD Ryzen 7 3700X, 64GB DDR4 RAM, Nvidia RTX 2060.

### Electrode localization

Electrodes’ locations were registered in the three-dimensional Montreal Neurological Institute (MNI) coordinate space (Evans et al., 1993). Electrodes were localized through coregistration of an individual subject’s T1 MRI scan with their CT scan using the Python package img_pipe (Hamilton et al., 2017). Three-dimensional reconstructions of the pial surface were created using an individual subject’s T1 MRI scan in Freesurfer and anatomical regions of interest for each electrode were labeled using the Destrieux parcellation atlas (Dale et al., 1999; Destrieux et al., 2010). These reconstructions were then inflated for better visualization of intra-Sylvian structures such as the insula and Heschl’s gyrus via Freesurfer. To visualize electrodes on the new inflated mesh, electrodes were projected to the surface vertices of the inflated mesh, which maintained the same number of vertices as the default pial reconstruction. To preserve electrode location using inflated visualization, each electrode was projected to a mesh of its individual Freesurfer ROI before projection to inflated space. Additionally, any depth electrodes greater than 4 millimeters from the cortical surface (*n* = 691) were not visualized on inflated surfaces due to a previously identified spatial falloff in high gamma frequency bands for electrodes greater than 4 millimeters apart from each other (Muller et al., 2016). Electrodes greater than 4 millimeters from the cortical surface, while excluded from visualization, were included in analyses if they contained a robust response (*p* < 0.05 for bootstrap procedure, *r* ≥ 0.1 for TRF modeling) to any task stimuli. To visualize electrodes across subjects, electrodes were nonlinearly warped to the cvs_avg35_inMNI152 template reconstruction (Dale et al., 1999) using procedures detailed in (Hamilton et al., 2017). While nonlinear warping ensures individual electrodes remain in the same anatomical region of interest as they were in native space, it does not preserve the geometry of individual devices (depth electrodes or grids). For inflated visualization in warped space, an identical ROI-mesh-to-inflated-surface projection method as described above was utilized, but the ROI and inflated meshes were generated from the template brain instead. Anatomical regions of interest were always derived from the electrodes in the original participant’s native space.

### Overt reading and playback task

#### Stimuli and procedure

The task was designed using a dual perception-production block paradigm, where trials consisted of a dyad of sentence production followed by sentence perception. Both perception and production trials were preceded by a fixation cross and broadband click tone (Figure 1A). Production trials consisted of participants overtly reading a sentence, then the trial dyad was completed by participants listening to a recording of themselves reading that produced sentence. Playback of this recording was divided into two blocks of consistent and inconsistent perceptual stimuli: consistent playback matched the immediately preceding production trial, while inconsistent playback stimuli were instead randomly selected from the previous block’s production trials. The generation of perception trials from the production aspect of the task allowed stimulus acoustics to be functionally identical across conditions.

Sentences were taken from the MultiCHannel Articulatory (MOCHA) database, a corpus of 460 sentences that include a wide distribution of phonemes and phonological processes typically found in spoken English (Wrench, 1999). A subset of 100 sentences from MOCHA were chosen at random for the stimuli in the present study; however, before random selection, 61 sentences were manually removed for either containing offensive semantic content or being difficult for an average reader to produce to reduce extraneous cognitive effects and error production, respectively. This task is identical to the one used in (Kurteff et al., 2023); see that paper for an analysis of this task in noninvasive scalp EEG.

For this study, a modified version of the task optimized for participants with a lower reading level was created so that pediatric participants could perform the task as close to errorless as possible. This version took the randomly selected MOCHA sentences from the main task and shortened the length and utilized higher-frequency vocabulary that still encompassed the range of phonemes and phonological processes found in the initial dataset. Seven of the seventeen participants (TC1, TC3, DC10, DC12, DC13, DC16, DC17) completed the easy-reading version of the task. Participants completed the task in blocks of 20 sentences (25 sentences for the easy-reading version) produced and subsequently perceived for a total of 40 (50) trials per block. Participants produced (and listened to subsequent playback of) an average of 142±61 trials. A broadband click tone was played in between trials.

Stimuli were presented in the participant’s hospital room on Apple iPad Air 2 using custom interactive software developed in Swift (Apple). Auditory stimuli were presented at a comfortable listening level via external speakers. Insert earbuds and/or other methods of sound attenuation (e.g., soundproofing) were not possible given the clinical constraints of the participant population. Visual stimuli were presented in a white font on a black background after a 1000 msec fixation cross. Accurate stimulus presentation timing was controlled by synchronizing events to the refresh rate of the screen. The iPad was placed on an overbed table and trials were advanced by the researcher using a Bluetooth keyboard. Participants were instructed to complete the task at a comfortable pace and were familiarized with the task before recording began. Timing information was collected by an automatically generated log file to assist in data processing.

#### Electrode selection

As mentioned above, electrodes >4 millimeters from the cortical surface were automatically excluded from visualization. However, electrodes identified as outside the brain or its pial surface via manual inspection of the subject’s native imaging were excluded from all analyses. Electrodes in a ventricle or in a lesion were excluded using the same method. Adjacent electrodes that displayed a similar response profile to outside-brain electrodes were also excluded; conversely, electrodes on the lateral end of a device that displayed a markedly different response profile than medially adjacent electrodes were determined to be outside the brain and thus excluded. As an additional measure of manual artifact rejection, channels that displayed high trial-to-trial variability were excluded from analysis. Lastly, while data were common average referenced in analysis, the data were re-preprocessed using a bipolar reference and any electrodes with a markedly different response when the referencing method was changed were excluded from analysis. All electrodes rejected through manual inspection of imaging were discussed and agreed upon by three of the authors (GLK, AF, LSH). Electrodes above the significance threshold (*p* > 0.05) for both perception and production, as determined by bootstrap procedure described below, were excluded from cNMF clustering if the electrode also had a low correlation during the mTRF modeling procedure (*r* < 0.1). In other words: electrodes without a significant perception or production response to sentence onset nor a moderate performance during mTRF model fitting were excluded from cNMF.

### Speech motor control task

#### Stimuli and procedure

A subset of six participants (TC6, DC7, DC10, DC13, DC16, DC17) completed a supplementary task with the goal of obtaining nonspeech oral motor movements to use as a control comparison for any electrodes that were production-selective to determine if they were speech-specific or not. Stimuli for this task consisted of written instructions accompanying a “go” signal on the iPad screen to prompt the participant to follow the instructions. The nine possible instructions, presented in a random order, were: “smile,” “puff your cheeks,” “open and close your mouth,” “stick your tongue out,” “move your tongue left and right,” “tongue up (tongue to nose),” “tongue down (tongue to chin),” and “say ‘aaaa,’” “say ‘oo-ee-oo-ee.’” These instructions were chosen as a subset of movements evaluated during typical oral mechanism exams conducted by speech-language pathologists (St. Louis & Ruscello, 1981). Each movement was repeated 3 times.

#### ERP analysis

For the nonspeech oral motor control task, except for the last two instructions (say “aa” or “oo-ee-oo-ee”), oral motor movements did not include an acoustic component. Thus, instead of being epoched to the acoustic onset of the trial like the primary task, responses were instead epoched to the display of the instruction text before the “go” signal, which was accompanied by the same broadband click tone as the main task. A match filter, identical to the one described above used to align high-resolution task audio with clinical recordings, identified the timing of these clicks and assisted in generation of the event files.

## Quantification and statistical analysis

### Event-related potential (ERP) analysis

We annotated accurate timing information for words, phonemes, and sentences to epoch data to differing levels of linguistic representation. A modified version of the Penn Phonetics Forced Aligner (Yuan & Liberman, 2008) was used to automatically generate Praat TextGrids (Boersma & Weenink, 2013) using a transcript generated by the iPad log file. Automatically generated TextGrids were checked for accuracy by the first author (GLK). Event files containing start and stop times for each phoneme, word, and sentence, as well as information about trial type (perception vs. production), were created using the iPad log file and accuracy-checked TextGrids. These event files were then used to average Z-scored high gamma across trials relative to sentence onset. For both production and perception, the onset of the sentence was treated as the acoustic onset of the first phoneme in the sentence as identified from the spectrogram. Responses were epoched between -0.5 and +2.0 seconds relative to sentence onset, with the negative window of interest intending to capture any pre-articulatory activity related to speech production (Chartier et al., 2018).

Electrode significance was determined by bootstrap *t*-test with 1000 iterations comparing activity during the stimulus to randomly selected inter-stimulus-interval activity; bootstrapped significance for perception and production activity were calculated separately as to identify electrodes that may be selectively responsive to either perceptual or production stimuli. For the bootstrap procedure, we averaged activity 5-550 milliseconds after sentence onset and compared that to average activity during a silent 400-600 milliseconds after the inter-trial click as a control. The control time window was selected as to not include potential evoked responses from the click sound but still be in the 1000 millisecond window between the click sound and stimulus presentation. A similar procedure was used to calculate significance for the consistent-inconsistent playback contrast (same time windows used). Bootstrap significance for the speech motor control task used activity 500-1000 milliseconds after the click sound played when text instructions were displayed to avoid including evoked responses to the click sound itself in the procedure. Because there were no inter-trial click sounds in the speech motor control task with the click instead marking the display of instructions, activity -500 to 0 milliseconds prior to the click sound was used as the control interval.

In addition to suppression, we were interested to see how onset responses change between speaking and listening. To quantify the presence of an onset response at a particular electrode, we looked in the first 300 msec of response relative to sentence onset for activity >1.5 SD above the mean response for the electrode’s activity epoched to sentence onset. The time window of the onset response was defined as the range of contiguous samples of activity >1.5 SD above the mean, with the peak amplitude of the onset response being the was greatest activity within the onset window. Onset latency was calculated as the maximum rate of change (differential) in the rising slope of the onset response. While we required an onset response to begin in the first 300 msec of activity after sentence onset, we did not specify a time window in which one must end. Onset responses were quantified separately for the average production response and average perception response of each electrode. Electrodes that exhibited an onset response during speech perception and production were classified as “dual onset,” while electrodes that exhibited an onset response during speech perception only were classified as “onset suppression.”

### Convex non-negative matrix factorization (cNMF)

To uncover patterns of evoked activity for speech production, speech perception, and auditory (click) perception that were consistent across participants, we employed convex non-negative matrix factorization (cNMF, Figure 3, (Ding et al., 2010)). This is an unsupervised clustering technique that reveals underlying statistical structure of datasets and has previously been used by our research group to discover profiles of neural response without explicitly specifying the feature represented by the response nor the anatomical location of the electrodes (Hamilton et al., 2018, 2021). We use a similar approach to these papers, summarized by the following equations:

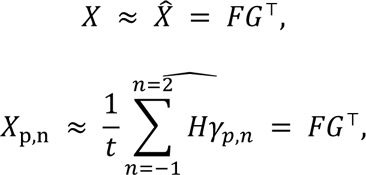

where *X* is the high gamma time series of shape (*n* samples, *p* electrodes) averaged across *t* epochs, and *F = XW*, where *W* is a matrix of shape (*p* electrodes, *k* clusters) and represents the cluster weights applied to the neural time series, and *G* is a matrix of shape (*p* electrodes, *k* clusters) and represents the weighting of an individual electrode within a cluster. cNMF was applied using this method to a concatenation of Z-scored evoked responses across subjects to sentences. Epochs consisted of a temporal range of -1 to +2 seconds relative to sentence onset. Epochs *t* were averaged within their response type then concatenated; possible response types were production onset, perception (playback) onset, and inter-trial click onset. Our method of performing cNMF on averaged epochs across different types of trials has been utilized in prior intracranial studies of speech (Leonard et al., 2019). In a supplemental analysis, we concatenated additional epoch averages corresponding to presentation of visual cues (e.g., text prior to reading, fixation cross) and a subdivision of playback onsets into consistent and inconsistent playback, but these manipulations did not significantly alter the clusters observed. We concatenated ERPs based on the response to production onset, perception (playback) onset, and click onset. We also incorporated information about expected vs. unexpected playback as well as presentation of the visual cue in separate supplemental analyses, but these did not significantly alter the clusters observed. Our final concatenation resulted in a matrix *X* of *n*\*3 samples (production epochs, perception epochs, click epochs) by *p* electrodes. The number of basis functions to include was determined by two primary factors: first, the identification of a threshold such that adding additional clusters resulted in diminishing increases in percent variance explained; second, identifying a point at which adding additional clusters resulted in redundant average responses across clusters. We calculated percent variance as the coefficient of determination (R^2^; Wright, 1921). This threshold was reached at *k*=9 clusters and 86% of the variance in the data explained. The average response for each of the *k*=9 clusters is provided in Figure S3.

### Suppression index (*SI*) calculation

Within the sentence-onset epochs, a further window of interest was defined to calculate the degree of suppression between task conditions. The window of interest for onset responses was defined as 0 to 1 seconds after sentence onset. Window sizes were determined by previous research on onset and sustained responses (Hamilton et al., 2018) as well as preliminary results of the unsupervised clustering technique shown in Figure 3. The suppression index (*SI*), or degree of suppression during speaking as compared to listening, was quantified at each electrode as the ratio of high gamma activity between two separate conditions averaged across all epochs for the task condition occurring at that electrode. This is formalized as:

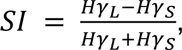

where *SI* of electrode *n* is the difference of high gamma activity during speaking (*HγS*) subtracted from high gamma activity during listening (*HγL*) divided by the sum of high gamma activity during speaking and listening in the first 1 second after the acoustic onset of the sentence. A positive *SI* means that activity was greater during listening as compared to speaking, whereas a negative *SI* means activity was greater during speaking compared to listening. An *SI* of zero would reflect no difference between conditions.

### Linear mixed-effects (LME) modeling

Linear mixed-effects (LME) models were fit using the package lmertest (Kuznetsova et al., 2017) in R at several points in analysis to quantify trends in the data. We chose LME as our statistical testing framework due to its ability to regress across within-and between-subject variability, facilitating generalization across subjects. The general equation takes the form:

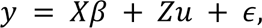

where *β* represents fixed-effects parameters, *u* represents random effects, and ε error. The first LME reported in this paper was used to quantify differences between suppression observed in onset and sustained responses. Suppression index (see above) was used as the response variable with window of interest (two-way categorical: onset or sustained) and ROI as fixed effects and subject as a random effect (in R: si ∼ window + roi + (1|subject)). *SI* was calculated separately in the onset and sustained windows for this analysis, unlike the *SI* calculation above: onset *SI* was calculated between 0 and 750 milliseconds and sustained *SI* was calculated between 1000 and 1750 milliseconds after sentence onset. We chose these windows based on the average duration of the onset response across all electrodes and chose to make the sustained time window non-contiguous with the onset window to prevent extraneous activity from longer onset responses erroneously being factored as sustained activity in the model. We reported the contrast in estimated marginal mean (EMM) *SI* of the two windows. We then used post-hoc Wilcoxon signed-rank tests with Benjamini-Yekutieli correction to calculate significant differences in *SI* between the onset and sustained responses within each ROI (Benjamini & Yekutieli, 2001). The second LME reported in this paper was used to quantify response latency within three regions of interest: primary auditory (HG, PT), non-primary auditory (STG, STS), and posterior + inferior insula. Peak latency values for the onset response (described above) were used as the response variable with ROI (three-way categorical) as a fixed effect and subject as a random effect (in R: peak_latency ∼ roi + (1|subject)). We reported the EMM peak latencies of the three ROIs as well as their contrasts. The third LME reported in this paper was used to quantify the mTRF ablation analysis, a causal probing technique where specific stimulus features are added or removed from an encoding model and differences in performance are recorded (Ivanova et al., 2021). For this LME model, the linear correlation coefficients between 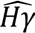 and *Hγ* were used as the response variable with model features (i.e., full vs. ablated) as a fixed effect and subject and channel as a random effect (in R: r ∼ model + (1|subject) + (1|channel)). We chose to include channel as a random effect here as we did not have a specific hypothesis for anatomical differences in ablated model performance; additionally, including channel as a fixed effect instead would have resulted in an uninterpretable amount of pairwise comparisons and introduce multiple comparisons bias into our analysis. We reported the EMM *r* values of the four models (base, ablate perception/production contrast, ablate consistent/inconsistent contrast, task-specific phonological feature encoding) as well as their contrasts. Contrast significance for all LMEs is calculated using *F* tests with Kenward-Roger approximation with *n* degrees of freedom specified, where *n* is the length of matrix *X* (Kenward & Roger, 1997).

### Multivariate temporal receptive field (mTRF) modeling

Multivariate temporal receptive field (mTRF) models were fit to describe the selectivity of the high gamma response to different sets of stimulus features (Aertsen & Johannesma, 1981; Crosse et al., 2016; Di Liberto et al., 2015; Theunissen et al., 2000). These models take the form of the equation below:

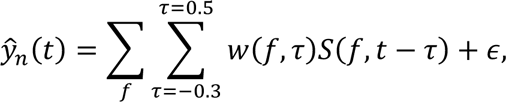

where 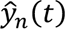 represents the estimated high gamma signal at electrode *n* at time *t*. The stimulus matrix *S* consists of behavioral information regarding features (*f*) for each time point *t* – τ, where τ is the time delay between the stimulus and neural activity. We fit separate models to predict the high gamma response in each channel using time delays of -0.3 sec to 0.5 sec. This delay range encompasses the temporal integration times to similar responses found in previous research (Hamilton et al., 2018), but with an added negative delay to encompass potential pre-articulatory neural activity (Chartier et al., 2018; Kurteff et al., 2023). Data were split 80-20 into training and validation sets. To avoid overfitting, the data were segmented along sentence boundaries, such that the training and validation sets would not contain information from the same sentence. These segments were then randomly combined into the 80/20 training/validation sets. Weights for each feature and time delay *w(f,τ)* were fit using ridge regression on the training set and a regularization parameter chosen by 10 bootstrap iterations. The ridge parameter was selected at the value that provided the highest average correlation performance across all bootstraps. Ridge parameters between 10^2^ and 10^8^ were tested in 20 logarithmically scaled intervals. Model performance was assessed using correlations between the high gamma response predicted by the model and the true high gamma response. Significance of these correlations was obtained through a bootstrap *t*-test procedure with 100 iterations in which the training data were shuffled in chunks to remove the relationship between the stimulus and response.

