## Supplemental Information for "Processing of auditory feedback in perisylvian and insular cortex"

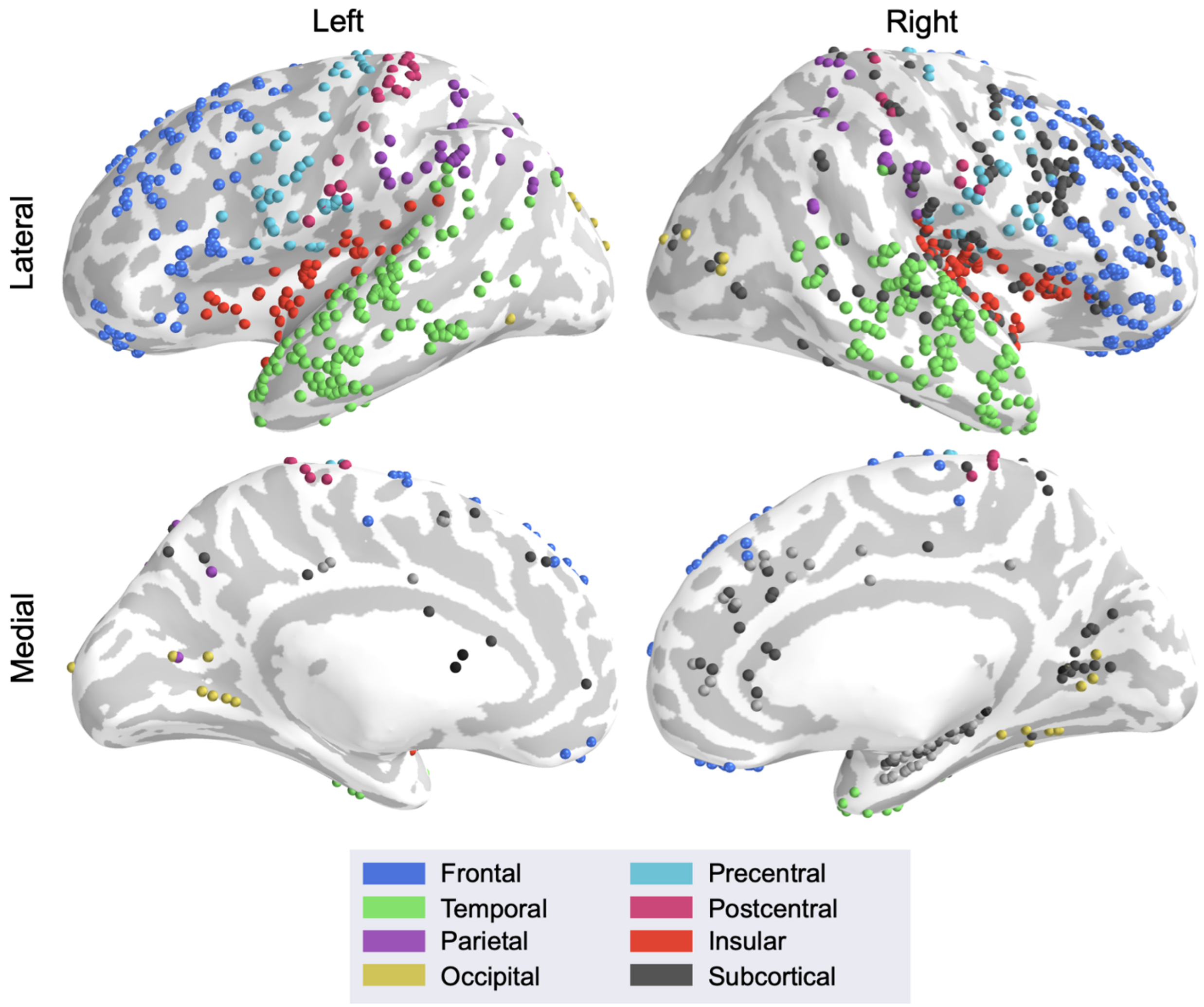


**Figure S1. Coverage map.**

Individual electrodes for all included subjects with imaging (*n* = 15; excluding TC1 & DC4) plotted on the cvs_avg35_inMNI152 atlas brain, color-coded by anatomical region of interest. Cortical surface inflated for better visualization of insular electrodes. Electrode visualization in native subject space is shown in Figure S2.

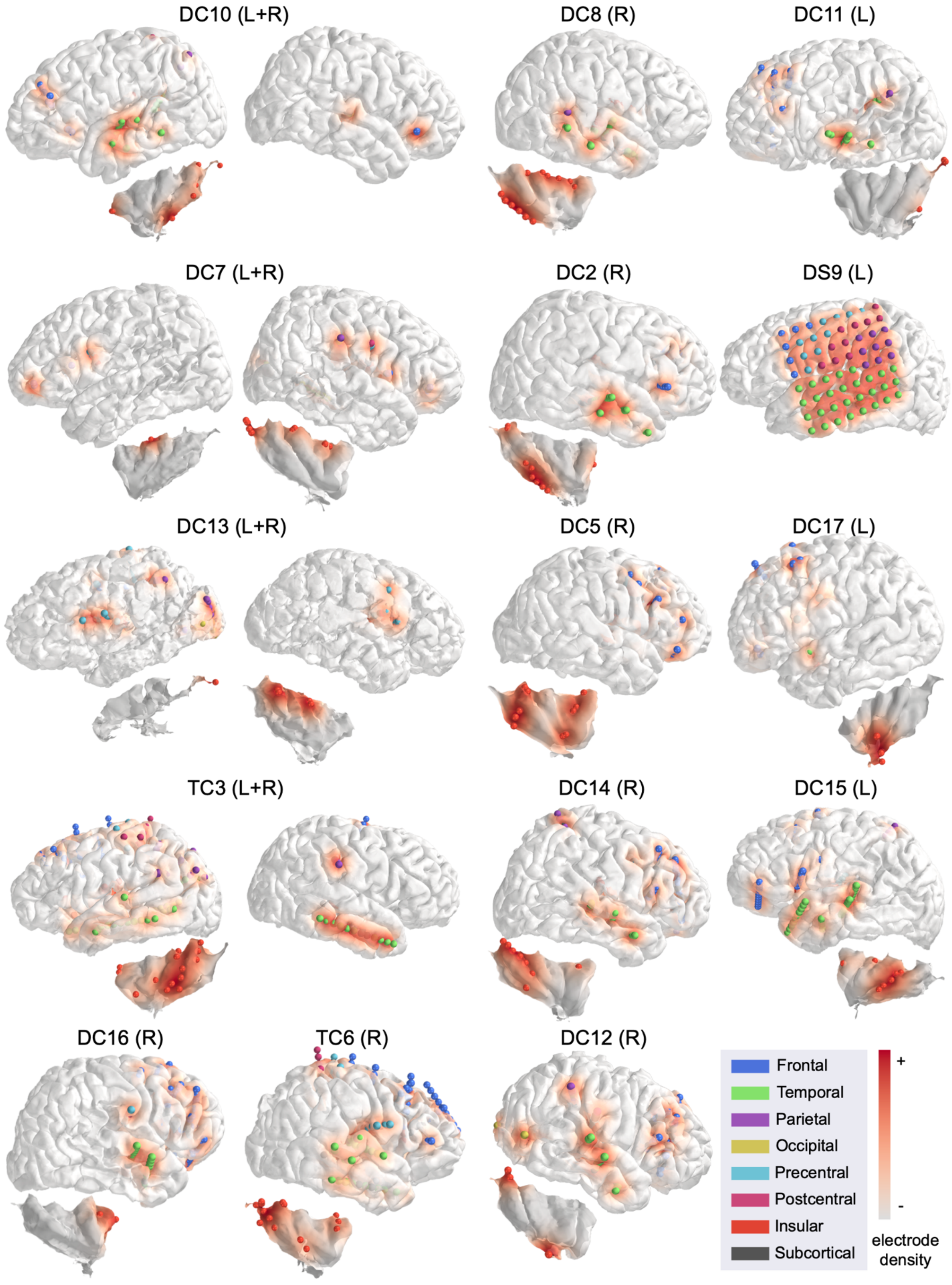

DC10 (L+R)
DC8 (R)
DC11 (L)
DC7 (L+R)
DC2 (R)
DS9 (L)
DC13 (L+R)
DC5 (R)
DC17 (L)
TC3 (L+R)
DC14 (R)
DC15 (L)
DC16 (R)
TC6 (R)
DC12 (R)
Frontal
Temporal
Parietal
Occipital
Precentral
Postcentral
Insular
Subcortical
+
-
electrode density

**Figure S2. Single subject coverage.**
Electrodes visualized on 3D reconstructions of individual subjects' MRIs, color-coded by anatomy. Color gradient represents density of electrode coverage. A separate reconstruction of individual subjects' insulas is provided for visualization of insular electrodes not visible from lateral cortical surface. Each subject displayed here is visualized on an averaged brain in Figure S1.

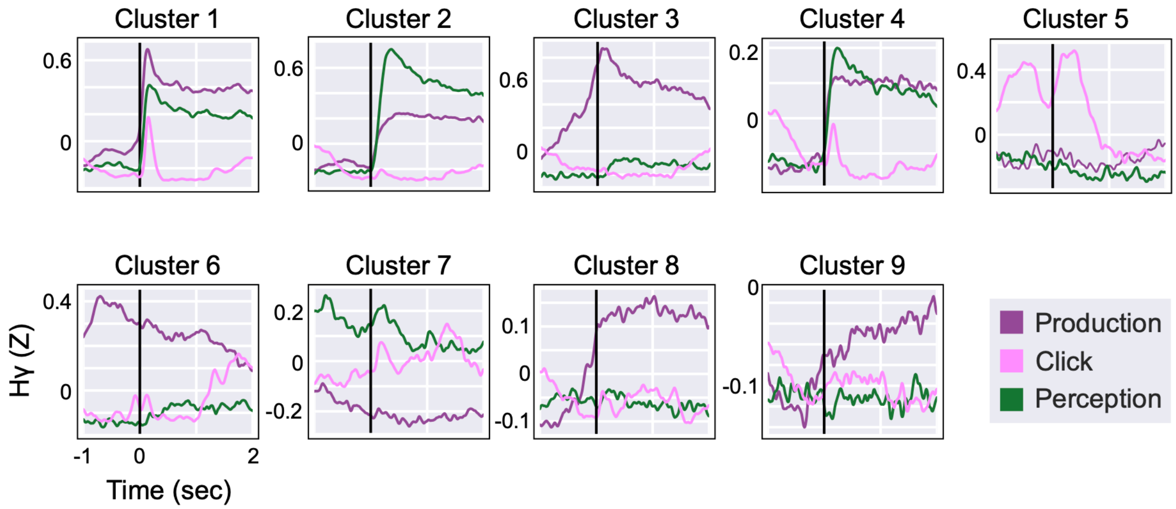


**Figure S3. Average response of all clusters in reported cNMF analysis.**
9 presented clusters explain 86% of the variance in the data (Figure 3A). "Onset Suppression" and "Dual Onset" clusters presented (Figure 3B) here are labeled as Clusters 2 and 1, respectively. "Pre-articulatory Motor" cluster presented (Figure 3B) here is labeled as Cluster 3. The responses plotted are the cluster basis functions of individual clusters relative to either sentence onset (production and perception conditions) or the inter-trial click tone (click condition).

| Participant | Age | Sex | Seizure focus |
|---|---|---|---|
| TC1 | 9 | M | Left temporo-parieto occipital |
| DC2 | 14 | M | Right hemisphere |
| TC3 | 19 | F | Left temporal |
| DC4 | 21 | M | No access to record |
| DC5 | 19 | M | Right frontal |
| TC6* | 14 | F | Right sensory frontal |
| DC7* | 20 | M | Right temporal |
| DC8 | 16 | F | No strong localization |
| DS9 | 37 | M | Left temporal |
| DC10* | 14 | X | Left temporal |
| DC11 | 13 | M | Bilateral frontal |
| DC12 | 14 | F | Right hemisphere and midline |
| DC13* | 8 | F | Left hemisphere |
| DC14 | 18 | F | Right frontal white matter |
| DC15 | 20 | F | Left frontotemporal |
| DC16* | 9 | F | Right frontal |
| DC7* | 17 | F | Left frontal |

**Table S1. Participant demographics.**
Table of age, sex, and seizure localization for each participant in this study. Subjects marked with an asterisk (*) completed the supplementary speech motor control task. Subjects in red were excluded from analysis due to a diagnosis of tuberous sclerosis complex. M = male, F = female, X = patient declined to

disclose. Letters in identifiers reflect recording site: TC = Texas Children’s Hospital; DC = Dell Children’s Medical Center; DS = Dell Seton Medical Center.
